# Renin Cells Drive Kidney Neurovascular Development and Arterial Remodeling when Renin Activity is Deficient

**DOI:** 10.1101/2025.07.08.663706

**Authors:** Manako Yamaguchi, Hiroki Yamaguchi, Jason P. Smith, Lucas Ferreira de Almeida, Daisuke Matsuoka, Alexandre G. Martini, Sara M Wilmsen, Sijie Hao, Kazuki Tainaka, Silvia Medrano, Maria Luisa S. Sequeira Lopez, R. Ariel Gomez

## Abstract

Renin cells synthesize and release the hormone-enzyme renin to regulate blood pressure and fluid-electrolyte homeostasis. Their function and identity depend on communication with surrounding cells and nerve fibers within complex kidney structure. Because renin cells are rare -0.01 % of kidney cells-conventional histological approaches cannot capture their interaction with nerve fibers and surrounding cells within the nephron and its vasculature. Using a novel ultrabright renin cell-specific tdTomato reporter mouse, high-resolution 3D imaging, and single-cell RNA-Seq, we mapped the interactions of renin cells with growing axons during normal kidney vascular development, in response to threats to homeostasis, and a severe arterial disease caused by a defective renin enzyme. During embryonic kidney development, stromal and renin cell progenitors assemble the arterioles, express axon attractants and neurotrophins that establish the precise innervation of renin cells and arterioles in a centrifugal pattern. Hypotension and sodium depletion led to an increase in the volume and number of renin cells along the arterioles. Renin enzymatic deficiency led to hypertrophy and endocrine transformation of renal arterioles, aberrant axon sprouting and sympathetic hyperinnervation suggesting a feed-forward mechanism whereby renin cells and axons co-induce each other, orchestrate neurovascular development and arteriolar remodeling when renin cells are over stimulated.

## Introduction

Renin cells appeared in nature over 400 million years ago, remaining essential for survival through evolution (1). They produce and secrete renin, a hormone that controls the key, initial step of the renin-angiotensin-system (RAS) to regulate blood pressure and fluid-electrolyte homeostasis (1, 2). In adult mammals, renin cells are localized to the juxtaglomerular (JG) apparatus at the distal end of renal afferent arterioles (AAs). Renin secretion from these cells is regulated by three main physiological mechanisms: (i) activation of β-adrenergic receptors by sympathetic nerves, (ii) signaling from macula densa cells in response to changes in tubular NaCl concentration, and (iii) direct sensing of blood pressure by the renin cell baroreceptors (2). Besides their role in maintaining homeostasis, renin cells are also essential precursors for proper development of the renal vasculature (1, 2). During early kidney development, renin-expressing cells are widely and heterogeneously distributed throughout the renal arterial tree. As the arteries mature, renin cells differentiate into smooth muscle cells (SMCs), pericytes, and mesangial cells (MCs), and the distribution of renin cells becomes restricted to the JG area in the adult kidney (3). The expression of renin throughout the developing renal vasculature is preserved as a cellular memory in renin cell descendants (4, 5). When homeostasis is threatened by hypotension, dehydration, sodium depletion, or treatment with RAS blockers, arteriolar SMCs dedifferentiate, recall the memory of the renin phenotype and reactivate renin production, a phenomenon known as “recruitment” (5–7). Whereas this plasticity works well in acute situations, constant stimulation of renin cells is detrimental to the kidney. For instance, chronic RAS inhibition due to long-term use of RAS inhibitors or inactivating mutations of any of the RAS genes causes concentric thickening of the intrarenal arterial tree: renin cells undergo phenotypic transformation into an embryonic-secretory invasive phenotype, which in turn induces SMCs to accumulate concentrically and attract sympathetic nerve fibers (2, 8–11). Over time, the arterioles are surrounded by inflammatory cells including macrophages and T cells (10, 11). Consequently, each renal arteriole is transformed into a neuro-immune-endocrine organ devoted mostly to the production of renin at the expense of kidney function and structural damage (10). These findings highlight the extraordinary plasticity of renin cells, which adapt their morphology, localization, and secretory profile in response to developmental cues and homeostatic challenges. However, traditional histological approaches using 2D tissue sections or microdissection of renal vessels have limitations, making it difficult to comprehensively capture dynamic changes while preserving structural relationships, including topological interactions with surrounding tissues and detailed patterns of innervation.

In this study, by combining advanced tissue clearing-based 3D imaging techniques with a newly developed mouse model that robustly labels renin-expressing cells (12), we visualized in great detail the dynamic changes in the distribution and innervation patterns of renin-producing cells from early developmental stages through adulthood, and ultimately under pathological conditions induced by chronic renin enzymatic insufficiency. Furthermore, by integrating single-cell RNA-Seq data from renal vascular mural cells, including renin cells, at multiple developmental stages, we uncovered stage-specific changes in the expression of molecules critical for guiding, establishing, and maintaining neurovascular units during development and disease. These findings provide fundamental insights into renin cells, their role in renal vascular development, organismal homeostasis, and vascular disease.

## Results

### Three-dimensional organization and distinct cellular architecture of renin cells within renal AAs

Using *Ren1^c-tdTomato/+^* mice (12), we show the precise distribution of individual renin cells and their relationship with mural cells within the 3D architecture of the renal arterial tree at single-cell resolution (Figure 1, A-D, Supplemental Video 1). Renin cells in adult mice were localized exclusively at the glomerular inlet near the terminal segments of the AAs (Figure 1, C and D).

**Figure 1.**
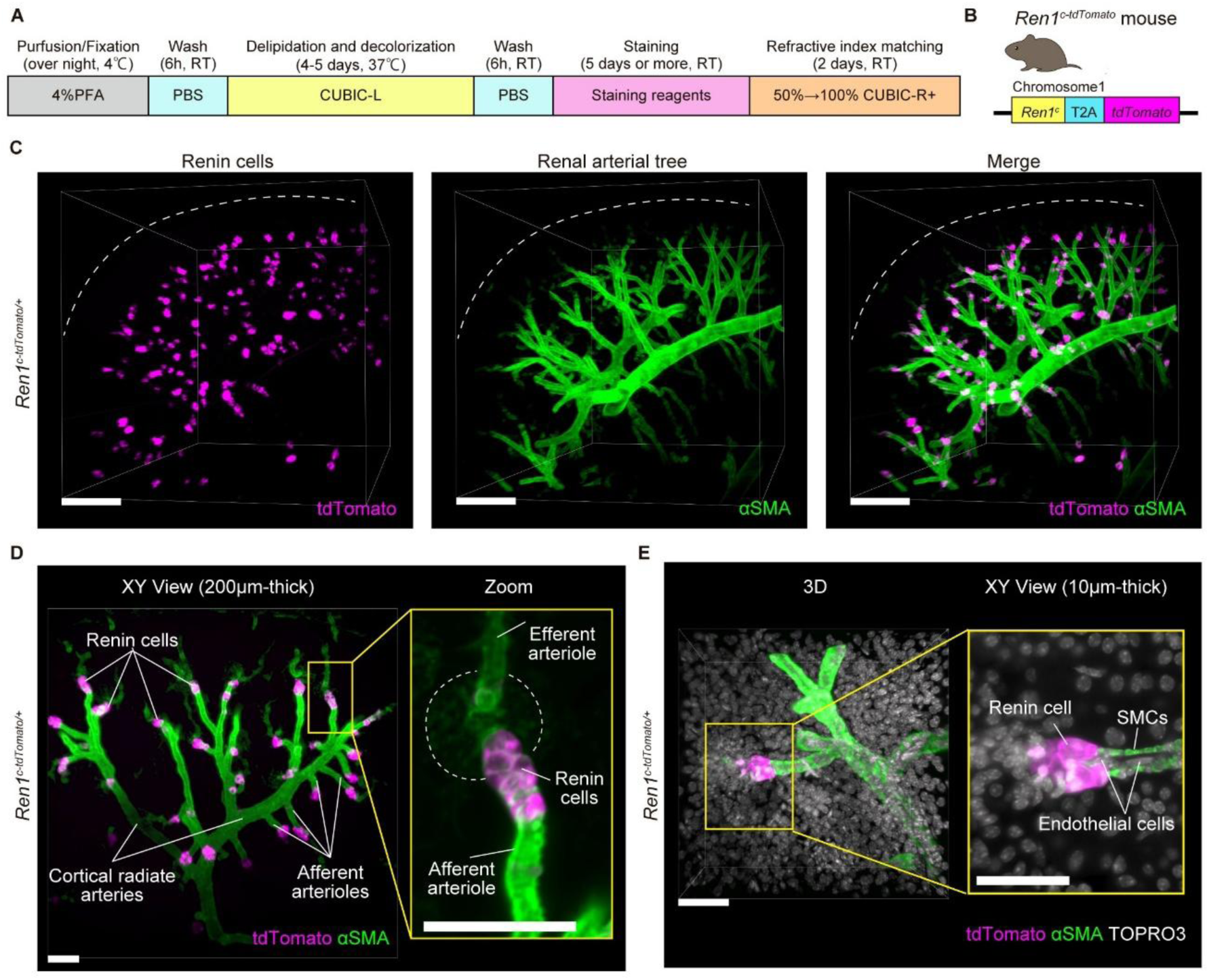
Tissue clearing-based 3D imaging of *Ren1^c-tdTomato/+^* mice allow high-resolution 3D structural observations of renin cells and vascular tree. (A) Workflow for kidney clearing and immunostaining (CUBIC protocol). (B) *Ren1^c-tdTomato^* mouse generation schematic. (C) Three-dimensional visualization of renin cells (tdTomato) and renal arterial tree within the kidney cortex of a 7-month-old *Ren1c^-tdTomato/+^* mouse. Dashed lines: kidney surface. Scale bars: 300 µm. (D) XY-plane (200 µm thickness) showing detailed structure of peripheral renal arterial tree (green) and localization of renin cells. Right panel: cell-level detail. Dashed circle: glomeruli. Scale bars: 100 µm. (E) High-resolution 3D image depicting spatial relationships between renin cells, smooth muscle cells (SMCs), and nuclei in the kidney cortex of a *Ren1^c-tdTomato/+^* mouse. Right panel: Enlarged XY-plane view (10 µm thickness) illustrating spatial relationships among renin cells, endothelial cells, and SMCs. Scale bars: 50 µm. RT, room temperature; αSMA, α-smooth muscle actin. See also Supplemental Videos 1 and 2.

Staining with TOPRO3 showed nuclei, cellular shape and spatial arrangement of vascular SMCs, renin cells, and their relationship with endothelial cells (Figure 1E, Supplemental Video 2). In renal arterioles, endothelial cells were oriented horizontally, aligning their nuclei along the longitudinal axis of the blood vessel lumen. In contrast, SMCs formed a single layer arranged perpendicularly around the endothelial layer, constituting the mural vessel wall. Although continuous with SMCs, renin cells exhibited a distinct rounded morphology, with their nuclei positioned eccentrically toward the cell periphery. Renin cells formed characteristic raspberry-like clusters, and endothelial cells lined the luminal surface of these clusters. Notably, endothelial cells were less densely packed compared to SMCs and renin cells, potentially facilitating efficient transmission of blood flow- and pressure-related mechanical signals from individual endothelial cells to multiple surrounding SMCs and renin cells.

### During physiological stress, renin cells swell and increase in numbers and SMCs along the AAs switch to the renin phenotype

Under physiological stress, such as volume depletion, hypotension or inhibition of the RAS, cells along renal arterioles are recruited to synthesize and release renin (1, 2). To visualize changes in the distribution of renin cells following a short-term physiological stress, we treated a group of 3-month-old *Ren1^c-tdTomato/+^* mice with captopril and low sodium diet for eight days (5, 13). The intensity of tdTomato fluorescence in renin cells increased significantly, and renin cell clusters enlarged in comparison to controls (Figure 2A). To quantitate these changes, we extracted a total of 121 control and 129 treated AAs from the reconstructed renal arterial trees from two mice per group. In each arteriole, we confirmed 3D continuity from the arcuate artery to the glomerulus and measured the continuous length of the renin-expressing segment (renin cluster length) extending from the glomerulus upstream the arteriole (Figure 2B). The histogram in Figure 2C clearly illustrates that the renin cluster length increased upon treatment. Indeed, the median renin cluster length was significantly longer in treated animals compared to controls (Supplemental Figure 1A). We further classified the AAs into cortical and juxtamedullary groups based on their branching location (Figure 2B). Under control conditions, cortical AAs exhibited significantly longer renin cluster lengths compared to juxtamedullary AAs (*P* = 0.001), but no significant difference was observed between these two groups after short-term treatment with captopril and low sodium diet (Figure 2D).

**Figure 2.**
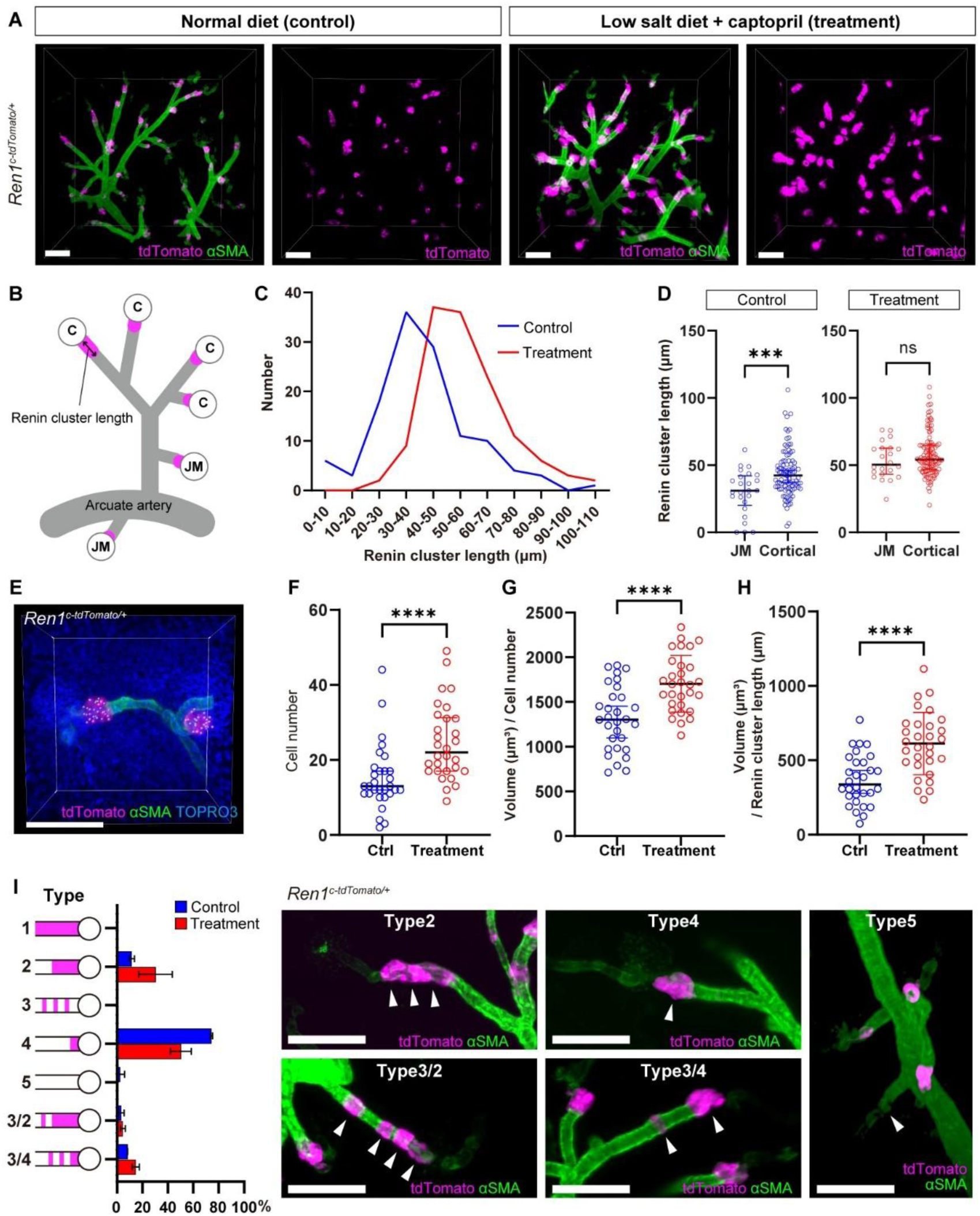
Acute physiological stress significantly alters the number, volume, and distribution of renin cells in afferent arterioles (AAs). (A) Representative 3D images of kidney cortex from 3-month-old *Ren1^c-tdTomato/+^* mice (control vs. low-salt diet + captopril). Under physiological stress, renin cells exhibited enhanced brightness, swelled in juxtaglomerular (JG) regions, and extended upstream along the AAs, indicative of recruitment. Scale: 100 µm. (B) Schematic of cortical (C) and juxtamedullary (JM) AA structure. Renin cluster length was defined as the longitudinal extension of renin cells along the AA from the glomerular inlet. (C) Histogram showing elongated renin cluster lengths under physiological stress. (D) Renin clusters were shorter in JM-AAs than cortical AAs in controls (*P*=0.0009), but not under physiological stress (*P*=0.3287). Data from 121 control and 129 treated AAs (2 mice/group; Mann–Whitney U test). (E) 3D reconstruction of renin cell nuclei (Imaris spot). Scale bar: 100 µm. (F–H) Cell number (F), cluster volume per cell (G), and volume per length (H) significantly increased by treatment. Data from 30 AAs/group (2 mice/group; Mann– Whitney U test (F), Student’s t-test (G, H)). (I) Left panel: Classification and frequency of renin localization types. Data represent measurements from two mice per group. Right panels: Representative 3D images illustrating each type. White arrowheads indicate renin cell localization patterns. Scale bars: 100 µm. ***, *P* < 0.001; ****, *P* < 0.0001; αSMA, α-smooth muscle actin; ns, no significant difference; Ctrl, control.

Next, to determine whether the enlargement of renin clusters was due to an increase in cell number or cellular hypertrophy, we performed 3D nuclear staining analyses (Figure 2E). We examined 30 control and 30 treated renin clusters from cortical AAs and measured their volume, length, and the number of cells per cluster. The treatment significantly increased the number of renin cells compared to controls (Figure 2F). In addition, the ratios of cluster volume to cell number and cluster volume to cluster length were significantly higher in treated mice compared to controls (Figure 2, G and 2). These findings indicate that enlargement of each renin cluster in response to short-term physiological stress is due to increased cell number and individual cell hypertrophy. Moreover, in treated animals, numerous AAs exhibited a distinct recruitment pattern, characterized by tdTomato fluorescence extending longitudinally along the arteriole walls in a stripe-like manner, indicating newly recruited renin cells (Figure 2A). We classified the localization patterns of renin cells within AAs into seven previously defined types based on immunostaining data from micro-dissected rat renal arteries (3). Applying this classification to our dataset (n = 121 controls, n = 129 treated from two mice per group), we identified five types, excluding types 1 and 3 (Figure 2I). In control kidneys, 74.4% of AAs were classified as type 4, characterized by renin expression strictly confined to the JG region (Figure 2I). In contrast, under short-term physiological stress, the proportion of type 4 AAs decreased to 50.3%, accompanied by increases in other types indicative of expanded renin expression. For instance, Type 2, in which clusters of renin cells extend proximally from the glomeruli but do not cover the entire length, increased from 11.5% in the control group to 30.4% after treatment (Figure 2I). Type 3/2 or 3/4, a mixed type combining type 2 or 4 with a striped distribution pattern (type 3), increased from 11.5% to 19.3% (Figure 2I). In addition, type 5 arterioles (lacking renin expression) accounted for 2.5% in controls and were completely absent following treatment (Figure 2I). The renin-positive length of AAs, including the striped distribution pattern, was significantly longer in the treatment group, with 14.7% exceeding 100 µm (Supplemental Figure1, B-D). In summary, detailed 3D images revealed morphological changes in renin cells induced by short-term physiological stress, including clear alterations in the size, cell number, and spatial distribution of renin cell clusters within the renal arterial tree.

### Sympathetic and sensory nerve fibers establish neuro-effector synapses with renin cells

To define how nerve fibers innervate renin cells, we performed 3D immunostaining with a fluorescently labeled anti-tubulin beta 3 (TUBB3) antibody, a pan-neuronal marker. In the adult *Ren1^c-tdTomato/+^* mouse kidney cortex, nerve fibers branched along the arterial tree, and upon reaching renin cells, emitted thin axonal branches that intimately intertwined with renin cells (Figure 3A, Supplemental Video 3). After making close contact with renin cells, axons continued to extend toward the glomerulus. To precisely visualize the 3D relationship between nerve fibers and the glomeruli, we also performed TUBB3 staining in *Ren1^c+/-^;Ren1^c-Cre^;R26R^mTmG^*, in which renin lineage cells are labeled with GFP and all other cells express tdTomato (8). The visceral layer of renal corpuscles strongly expressed tdTomato, clearly outlining the glomerular structure. Interestingly, after reaching the renin cells, axons did not penetrate into the glomerulus, where intraglomerular MCs reside. Instead, axons passed through the JG apparatus, extended along efferent arterioles, and further branched toward peripheral segments of the arterioles (Figure 3B, Supplemental Video 4). We next identified which types of axons innervated renin cells and the vasculature. Tyrosine hydroxylase (TH) immunostaining confirmed that the majority of fibers contacting renin cells were sympathetic nerve fibers (Figure 3C). Additionally, calcitonin gene-related peptide (CGRP)-positive thin sensory nerve fibers were also observed in close proximity to the renin cell clusters (Supplemental Figure 2). Furthermore, immunostaining with the synaptic marker synaptophysin confirmed the presence of synaptic vesicles along the nerve fibers associated with renin cells, indicating that “en passant” neuroeffector junctions are formed at these sites (Figure 3D). Consistent with these observations, electron microscopy revealed axon varicosities containing synaptic vesicles closely apposed to renin-secreting granular cells, further supporting the direct autonomic innervation of renin cells (Figure 3E).

**Figure 3.**
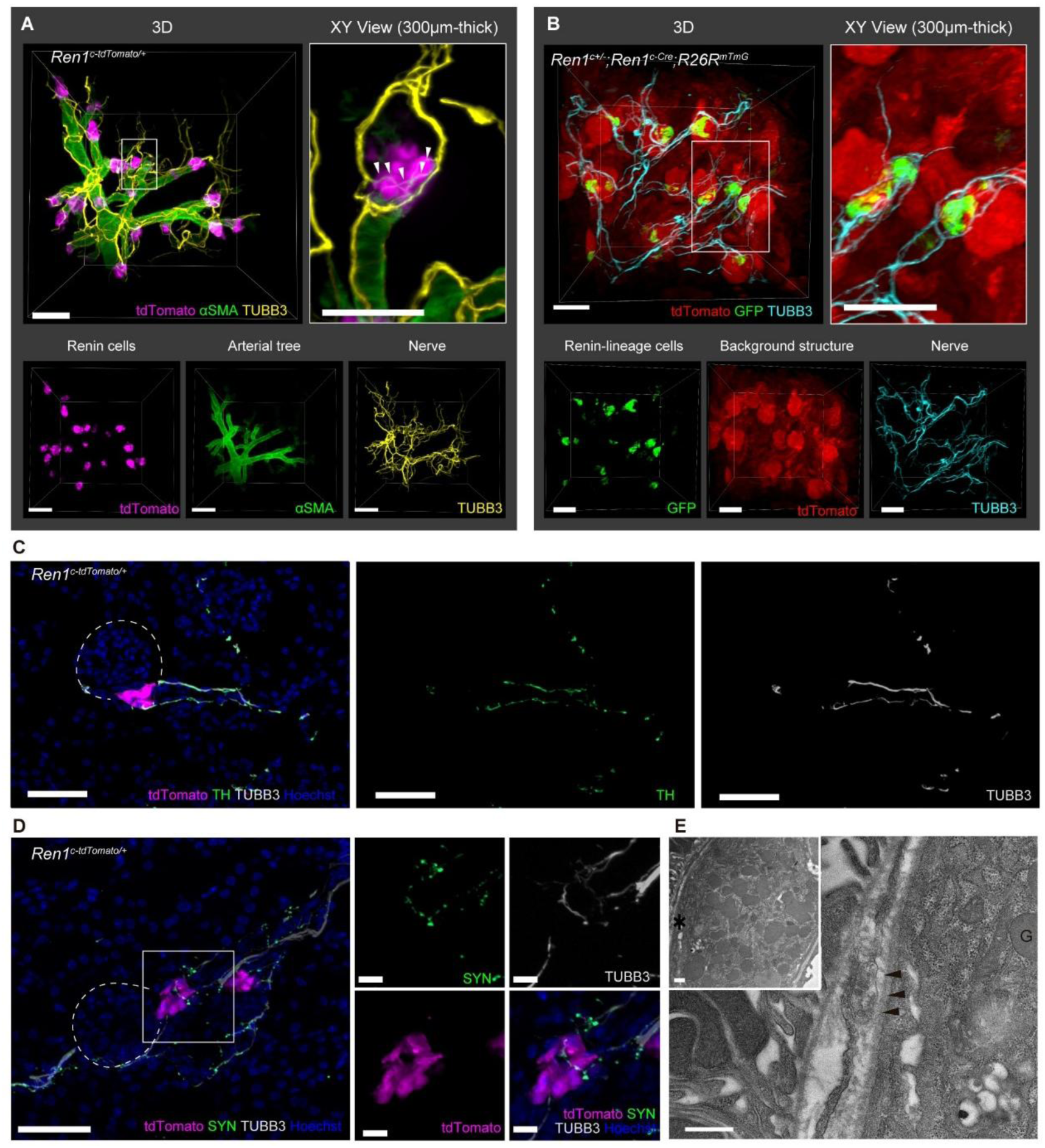
Dense sympathetic innervation of renin cell clusters in afferent arterioles. (A) 3D images of *Ren1^c-tdTomato/+^* mouse kidney cortex showing renin cells, arterial tree and nerve fibers. The enlarged XY view (right panel) highlights nerve fibers enveloping the renin cells, with fine branching of nerve fibers indicated by arrowheads. Bottom panels depict separate channels. Scale bars: left and bottom panels, 100 µm; enlarged view, 50 µm. (B) Three-dimensional images from *Ren1^c+/-^; Ren1^c-Cre^; R26R^mTmG^* mice labeled for renin-lineage cells (GFP), other tissue structures (tdTomato), and nerve fibers. Enlarged XY view highlights nerve fiber arrangement along afferent and efferent arterioles, avoiding glomeruli. Bottom panels depict separate channels. Scale bars: 100 µm. (C) Immunofluorescence staining of *Ren1^c-tdTomato/+^* mouse kidneys for tyrosine hydroxylase (TH) and tubrin-beta 3 (TUBB3), confirming sympathetic innervation of renin cells. Dashed circle: glomerulus. Scale: 50 µm. (D) Synaptic marker synaptophysin (SYN) co-localized with TUBB3-positive nerve fibers, suggesting synaptic connections with renin cells. Right panels: enlarged views showing colocalization. Dashed circle: glomerulus. Scale bars: left image, 50 µm; right enlarged panels, 10 µm. (E) Electron microscopy image illustrating nerve terminal containing synaptic vesicles (arrowhead), closely associated with a renin-secreting cell containing renin granules (G). Inset: low-magnification view of a renin cell. The main panel is an enlarged view of the region indicated by an asterisk (*). Scale bars: 500 nm. αSMA, α-smooth muscle actin. See also Supplemental Videos 3 and 4.

### During kidney development the time of appearance, morphology and direction of renin cells differ between juxtamedullary and cortical arterioles

During the development of the metanephric kidney, renin precursor cells originate from forkhead box protein D1 (FoxD1)-expressing stromal cells (2). These renin precursor cells subsequently differentiate into renin-producing JG cells, SMCs, MCs, and interstitial pericytes (14, 15). Renin precursor cells play a crucial role in the morphogenesis and branching of the developing renal arterial tree (15). In this study, we refer to cells expressing renin during kidney development as “renin-expressing cells”. Since mouse glomerulogenesis continues until approximately 7–10 days after birth, the neonatal renal cortex contains nephrons at various developmental stages (16). To define the developmental changes in the spatial distribution of renin-expressing cells, we performed whole-mount 3D imaging of developing kidneys at E18.5, P0, and P5 from *Ren1^c-tdTomato/+^* mice. At E18.5, renin-expressing cells were broadly distributed across the developing renal arterial tree, including both central and peripheral arterial segments. (Figure 4A). By P0, renin expression became more sporadic in cortico-medullary and arcuate arteries compared to E18.5; however, peripheral arteriole walls leading directly into the developing glomeruli remained densely covered by renin-expressing cells (Figure 4B, Supplemental Video 5). Notably, at E18.5 and P0, peripheral arterioles branching from arcuate arteries were predominantly oriented medially rather cortically, indicating early vascular growth toward juxtamedullary regions (Figure 4, A and B). In contrast, by P5, we clearly observed arterial branches extending from arcuate arteries toward the outer cortex. At this stage, further branching of peripheral arterial segments was evident, although arterial tree formation was not yet complete (Figure 4C). Renin-expressing cells were prominently localized in the JG regions, whereas their presence was more scattered along the peripheral arterioles extending from arcuate arteries toward the renal surface. Notably, at P5, renin cell clusters in the JG regions were larger and more conspicuous in juxtamedullary AAs compared to cortical AAs (Figure 4C, Supplemental Video 6). These results illustrate clearly the centrifugal pattern of arterial maturation during kidney development. In addition to confirming the conventional view that juxtamedullary glomeruli mature earlier than cortical glomeruli, our findings further suggest novel aspects of renal development: specifically, that the growth directions of AAs and the patterns of renin cell distribution differ between juxtamedullary and cortical arterioles during maturation.

**Figure 4.**
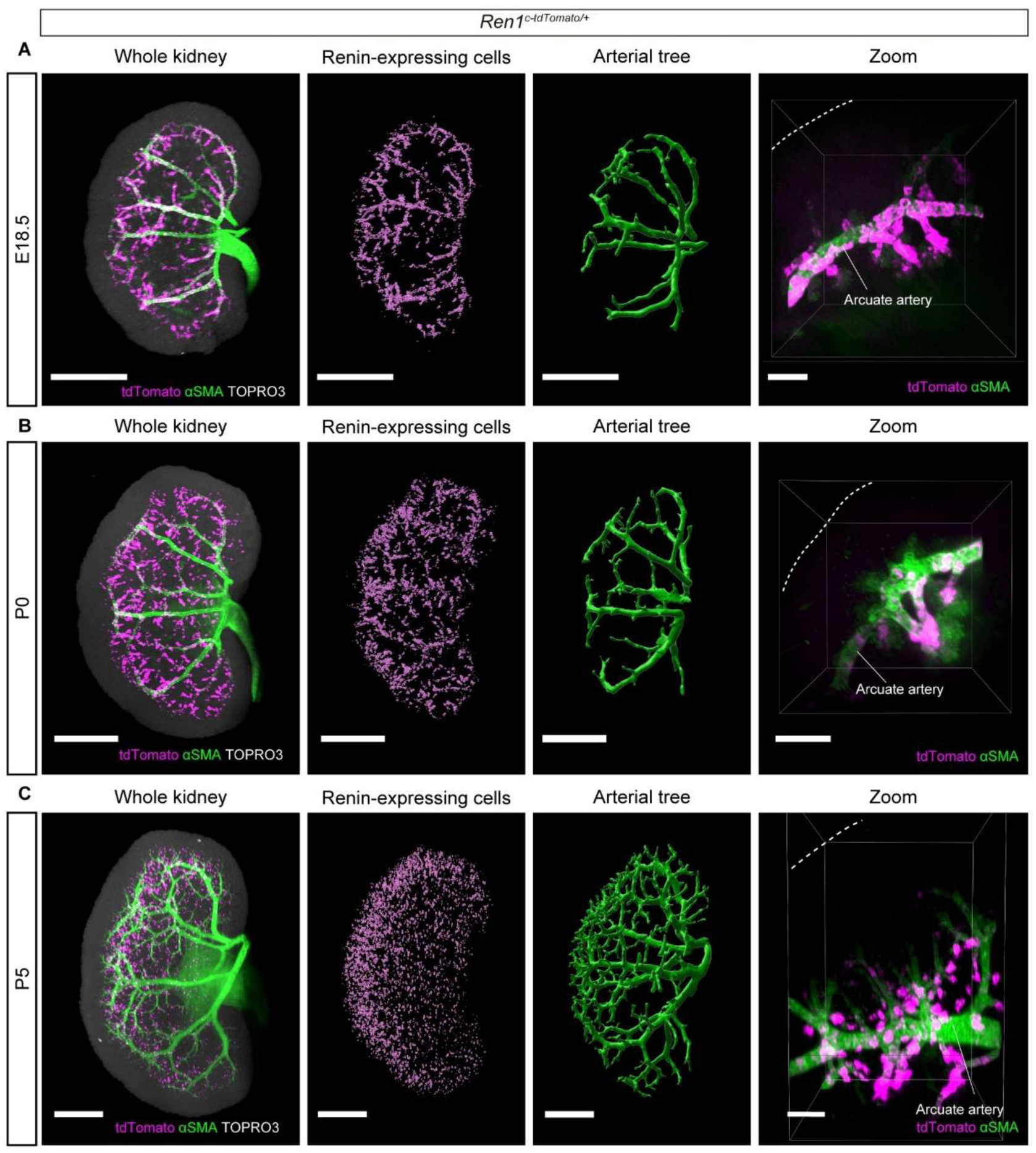
Distinct distribution patterns of renin-expressing cells during centrifugal maturation of renal arteries. (A–C) Representative whole-kidney 3D imaging of *Ren1^c-tdTomato/+^* mice at E18.5 (A), P0 (B), and P5(C). Panels show merged signals of renin-expressing cells, renal arterial tree and nuclei (left), extracted renin-expressing cells structure (second panel from left), and extracted arterial tree structure (third panel from left), and zoomed highlighting the spatial relationship between renin-expressing cells and the developing arterial tree, specifically at the arcuate artery level. Dashed lines: kidney surface. Scale bars: whole kidney and separated panels, 1 mm; zoomed panels, 100 µm. αSMA, α-smooth muscle actin. See also Supplemental Videos 5 and 6.

### Centrifugal maturation and innervation of the renal arterioles and renin cells during kidney development

Next, to gain deeper insight into the developmental pattern of innervation associated with centrifugal renal arterial maturation and renin cell differentiation, we examined kidneys from E17.5, P0, P5, P10, and 1-month-old *Ren1^c-tdTomato/+^* mice. Kidneys were subjected to 3D imaging with α-smooth muscle actin (αSMA) and TUBB3 staining. At E17.5, juxtamedullary AAs extending toward the medulla were clearly visible, and nerve fibers had already reached and become associated with renin-expressing cells along these arterioles. In contrast, within the arterial branches extending from the arcuate arteries towards the outer cortex, nerve fibers preceded the formation of αSMA-positive vascular walls. Furthermore, the tdTomato signals of renin-expressing cells in these cortical vessels appeared weaker compared to juxtamedullary AAs (Figure 5). Similar findings were observed at P0, with cortical arterioles starting to extend toward the outer cortex; however, tdTomato expression remained weak in αSMA-positive vascular walls, and nerve fibers preceded the maturation of the vascular wall structure (Figure 5, Supplemental Video 7).

**Figure 5.**
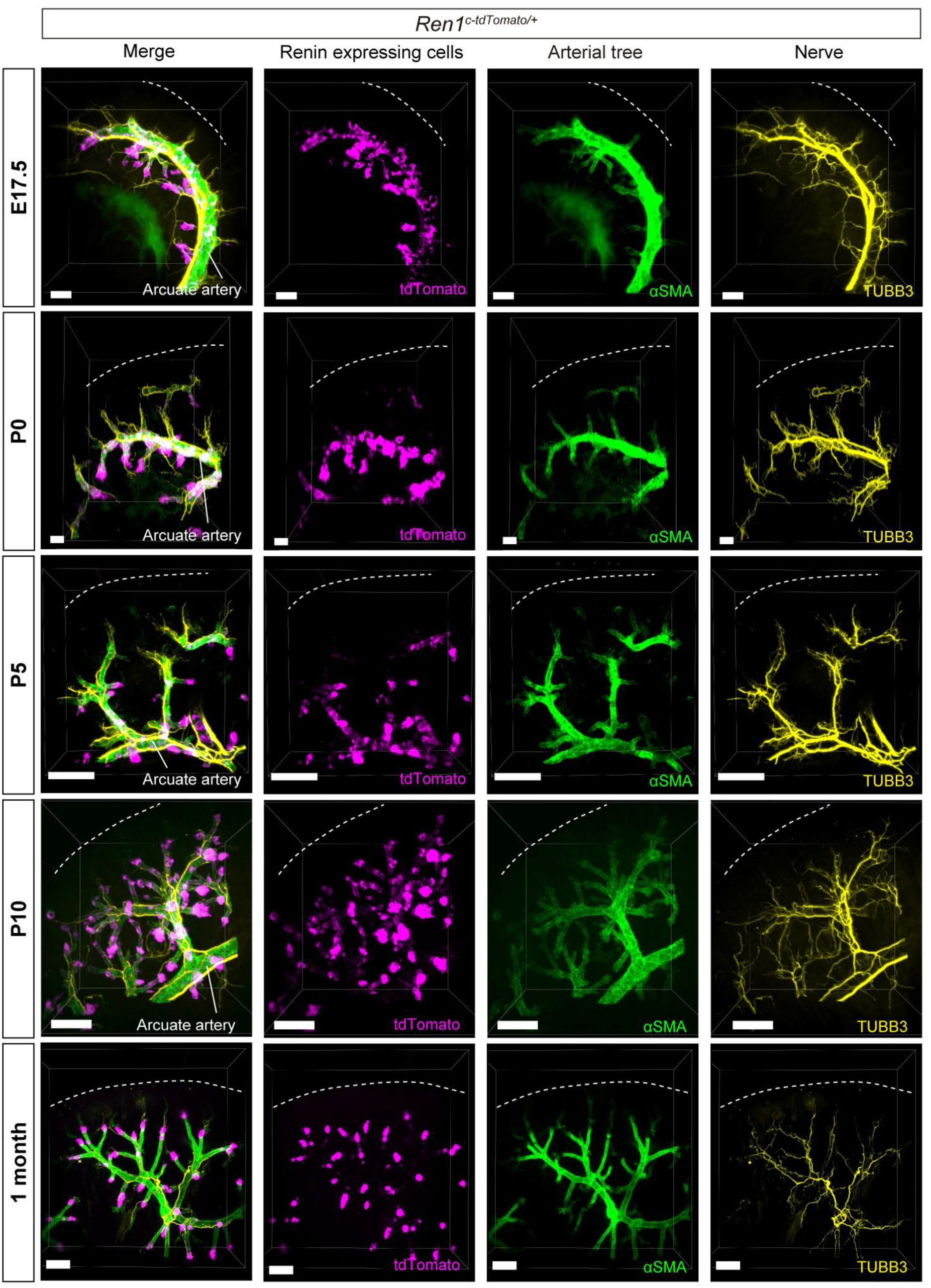
Nerve fiber development precedes renin cell differentiation in outer cortical regions. Three-dimansional views of renin cells, arterial tree, and nerves at E17.5, P0, P5, P10, and 1 month in *Ren1^c-tdTomato/+^* mice. At E17.5 and P0, juxtamedullary afferent arterioles extending medullary were covered by renin-expressing cells, whereas nerve fibers preceded cortical arterial wall. At P5, renin-expressing cell clusters appeared at cortical afferent arteriole tips connected to glomeruli and distributed in a striped pattern along the α-smooth muscle actin (αSMA)+ arterial tree. Peripheral nerve extension preceded renin-expressing cells. By P10, all arterial branches show fully innervated renin-expressing cells, becoming restricted to juxtaglomerular areas at 1 month. Dashed lines: kidney surface. Scale bars:: 100 µm. TUBB3, tubulin-beta 3. See also Supplemental Video 7.

By P5, prominent clusters of brightly fluorescent tdTomato-expressing cells were detected at the tips of cortical AAs that had completed their connection with developing glomeruli. Additionally, tdTomato-positive cells were arranged in a striped pattern along the αSMA-positive arterial walls; however, tdTomato expression remained weak in arterioles that were actively extending to the renal periphery. Notably, nerve fibers extended peripherally prior to the appearance of mature renin-expressing cells at this stage (Figure 5). At P10, renin-expressing cells were clearly observed at the terminal segments of all branches of the arterial tree, and each renin cell cluster was innervated. While tdTomato fluorescence was particularly intense within JG regions, a striped distribution pattern along arterioles remained visible (Figure 5). At 1 month, tdTomato expression became more localized specifically within the JG region (Figure 5). To confirm that tdTomato expression accurately reflects endogenous renin protein expression during developmental stages, we performed immunofluorescence staining and verified the precise colocalization of tdTomato signals with renin protein at both E17 and P5 stages (Supplemental Figure 3). Furthermore, 2D cross-sectional imaging confirmed that nerve fiber extension toward the renal outer cortex consistently preceded the appearance of renin-expressing cells (Supplemental Figure 3). In summary, our comprehensive 3D imaging analysis revealed distinct developmental dynamics in the relationship between nerve growth and the innervation and emergence of renin-expressing cells during the centrifugal maturation of the renal arterial tree.

### Foxd1^+^ progenitors, renin cells and ingrowing axons co-induce each other to build the neuro-arterial network of the developing kidney via neurotrophins and axon guidance molecules

As mentioned above, FoxD1-positive stromal cells are progenitors of renal vascular mural cells, including renin cells, SMCs, MCs, fibroblasts, and pericytes (17) (Supplemental Figure 4A). To gain further insights into the development of the arterial tree in the renal outer cortex and the distribution of renin-expressing cells, we performed 3D imaging analysis of kidneys from P5 *FoxD1^GC^; R26R^tdTomato^; Ren1^c-YFP^* mice. In this mouse model, cells derived from FoxD1-positive progenitors are labeled by tdTomato fluorescence, whereas cells actively expressing renin are marked by YFP fluorescence. The 3D imaging results revealed that blood vessels composed of FoxD1-lineage cells progressively extended toward the developing glomeruli in the outer cortex. In contrast, YFP-positive renin-expressing cells were exclusively observed in AAs connected to mature glomeruli (Figure 6A, Supplemental Video 8). Next, immunofluorescence staining further confirmed the positional relationship among FoxD1-lineage cells and renin, αSMA, TUBB3, and PECAM protein expression. Notably, nerve fibers marked by TUBB3 extended peripherally along the arterial wall composed of FoxD1-lineage cells, preceding the appearance of renin-expressing cells (Figure 6B). αSMA-positive cells were observed more peripherally than renin-positive cells, being present not only in renin-expressing regions but also extending further into peripheral, developing arterial segments lacking renin expression (Figure 6B). Nerve fibers were closely associated with αSMA-positive vascular regions (Figure 6B). Endothelial cells labeled by PECAM were already present within the arterial walls formed by FoxD1-lineage cells and in developing glomeruli at this developmental stage (Figure 6B).

**Figure 6.**
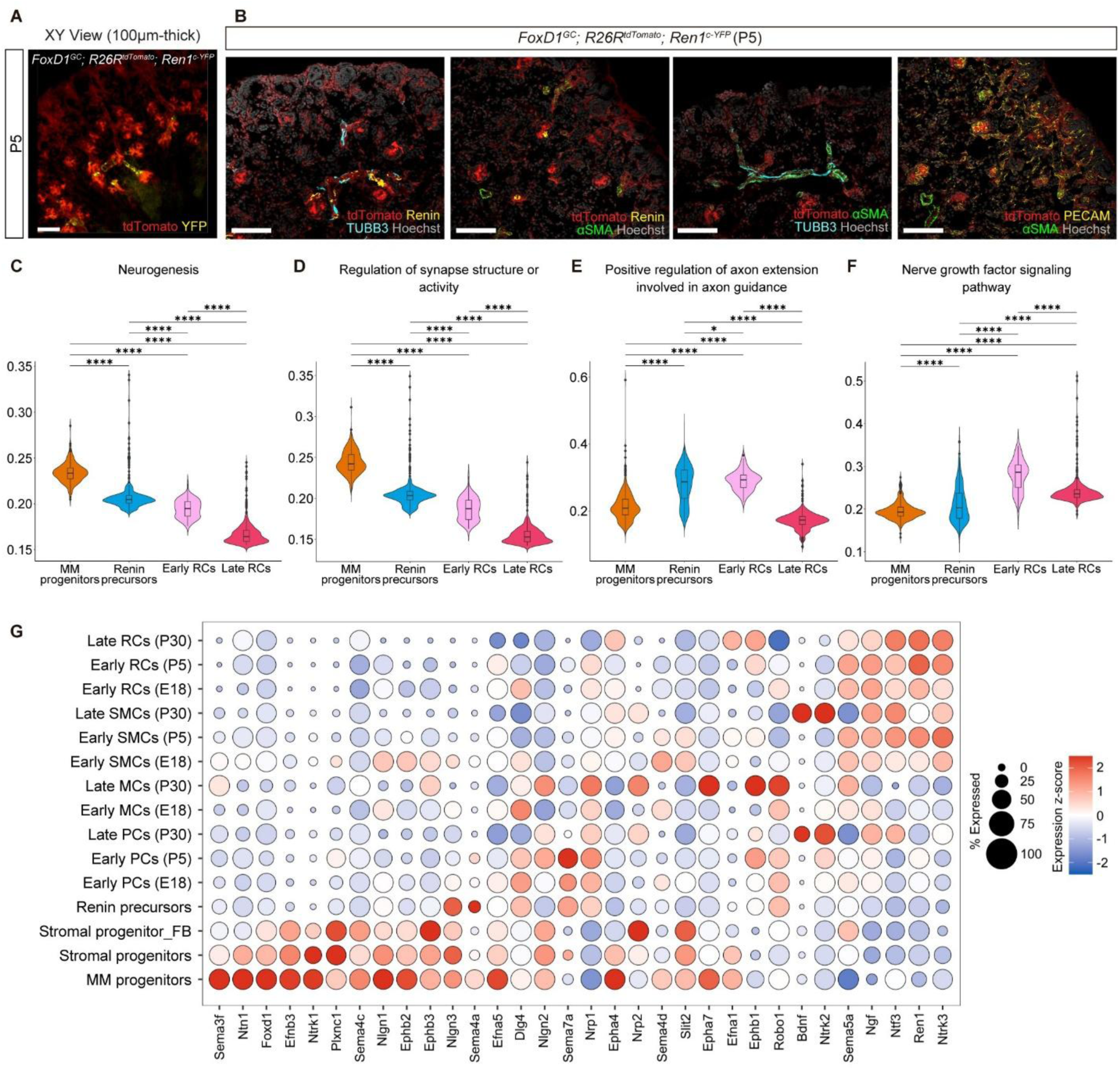
Stage-specific expression of axon guidance and neurotrophic factors by FoxD1-lineage cells in kidney development. (A) Representative XY-plane (100 µm thickness) 3D imaging of renal cortex from P5 *FoxD1^GC^; R26R^tdTomato^; Ren1^c-YFP^* mice, showing spatial distribution and relationship of renin-expressing cells (YFP) with FoxD1-lineage cells (tdTomato). Scale bar: 100 µm. (B) Immunofluorescence staining of P5 kidneys from *FoxD1^GC^; R26R^tdTomato^; Ren1^c-YFP^* mice illustrating detailed localization and interactions among renin cells, nerves, smooth muscle cells (SMCs) and endothelial cells. Scale bars: 100 µm. (C–F) Violin plots showing cell-specific single pathway analysis scores for neurogenesis (C), regulation of synapse structure or activity (D), positive regulation of axon extension involved in axon guidance (E), and nerve growth factor signaling pathway (F) across metanephric mesenchyme (MM) progenitors, renin precursors, early renin cells (RCs), and late RCs. Statistical comparisons were performed using pairwise t-tests with Benjamini-Hochberg multiple comparisons corrections; ****Padj < 0.0001, ***Padj < 0.001, *Padj < 0.05. (G) Dot plot showing stage-specific expression patterns of selected nerve-related genes and cell markers (*Foxd1* and *Ren1*) across different populations of FoxD1-derived renal cells at multiple developmental stages (E12, E18, P5, P30). Dot size: percentage of cells expressing the gene; color: expression levels (z-score). Data partly overlaps with Yamaguchi M et al., Circ Res, 2024, Figure C, though the visualization and cell categorization differ. αSMA, α-smooth muscle actin; TUBB3, tubulin-beta 3; MC, mesangial cell; PC, pericyte; FB, fibroblast. See also Supplemental Video 8.

To elucidate how FoxD1-lineage cells participate in nerve-related signaling throughout kidney development, we utilized our previously published single-cell RNA-Seq dataset (GSE218570), derived from kidneys of *FoxD1^GC^; R26R^mTmG^* mice at E12, E18, P5, and P30 (4, 18) (Supplementary Figure 4, B and C). Single Pathway analysis in Single Cells (SiPSiC) (19) was applied, and pathway activation scores were visualized across distinct cell populations, including metanephric mesenchyme (MM) progenitors, renin precursors, early renin cells, and late renin cells (Figure 6, C-F). Gene ontology (GO) terms associated with nerve development demonstrated clear stage-dependent expression patterns. Genes involved in neurogenesis and the regulation of synapse structure or activity were predominantly enriched in earlier developmental populations, specifically MM progenitors and renin precursors (Figure 6C, 6D). Genes related to the positive regulation of axon extensions involved in axon guidance were notably enriched during intermediate developmental stages, particularly in renin precursors and early renin cells (Figure 6E). Meanwhile, genes associated with the nerve growth factor (Ngf) signaling pathway increased in expression during later stages of differentiation, with enrichment becoming prominent in early renin cells, and remaining elevated in the late renin cell stage (Figure 6F). Next, we visualized nerve-related genes identified in multiple GO terms, along with *Foxd1* and *Ren1* markers, in a dot plot to characterize their expression during FoxD1-lineage cell differentiation (Figure 6G). During kidney development, these nerve-related genes exhibited distinct, characteristic, and stage-specific expression patterns. Undifferentiated FoxD1-derived progenitor cells strongly expressed axon guidance cues, including semaphorin 3F (*Sema3f*), netrin-1 (*Ntn1*), ephrin-B3 (*Efnb3*), plexin C1 (*Plxnc1*), semaphorin 4C (*Sema4c*), neuroligin-1 (*Nlgn1*), ephrin type-B receptor 2 (*Ephb2*), ephrin type-B receptor 3 (*Ephb3*), neuroligin-3 (*Nlgn3*), semaphorin 4A (*Sema4a*), and ephrin-A5 (*Efna5*), suggesting their role in guiding nerve fibers to precise locations within the developing renal tissue. Interestingly, certain axon guidance genes, notably *Efna1, Epha4*, and *Ntn1,* exhibited a biphasic expression pattern with peaks at both the MM progenitor and late renin cell stages (Figure 6G, Supplementary Figure 4D). This biphasic expression suggests dual functional roles, initially in cellular positioning and nerve fiber induction, and subsequently in cell differentiation and functional maturation. Renin precursor cells were characterized by high expression of semaphorin 4A (*Sema4a*), known as a neuroimmune semaphorin (20). Moreover, differentiated renin cells and SMCs showed elevated expression of neurotrophins, such as nerve growth factor (*Ngf),* neurotrophin-3 (*Ntf3*), and brain-derived neurotrophic factor (*Bdnf),* suggesting their role in stabilizing and maintaining innervation. Intriguingly, *Ngf* displayed an inverse expression pattern compared to its receptor neurotrophic receptor tyrosine kinase 1 (*Ntrk1)*, implying that Ngf acts primarily as a paracrine signal toward adjacent nerves (Figure 6G, Supplementary Figure 4D). Conversely, the expressions of *Bdnf* and its receptor *Ntrk2,* and *Ntf3* and its receptor *Ntrk3*, were closely correlated, indicating potential autocrine signaling roles or interactions between nerves and renin cells or SMCs (Figure 6G, Supplementary Figure 4D). In summary, FoxD1-lineage cells sequentially contribute to kidney neural network formation, transitioning from roles in early neurogenesis and axon guidance during initial differentiation, to stabilization and maintenance of neural networks during later maturation stages.

### Renin enzymatic insufficiency leads to overstimulation of renin cells, Ngf synthesis and aberrant vascular hyperinnervation

Genomic or pharmacological ablation of the RAS leads to a severe form of renal arterial disease characterized by the progressive thickening of the renal arterial tree (8, 9, 11). However, no physiological models existed to explore defects on the function of the renin protein itself. To address this, we generated a novel mouse model where the renin protein is synthesized but its enzymatic activity is reduced. The *Ren1^c-tdTomato^* mouse line we generated and used in this study has a normal phenotype in the hemizygous state (*Ren1^c-tdTomato/+^*). However, homozygous mice (*Ren1^c-tdTomato/tdTomato^*) have impaired renin enzymatic activity, leading to reduced angiotensin generation even when they produce and release high levels of circulating renin (12). Thus, these renin cells -with their blood pressure and volume sensing mechanisms otherwise intact-produce and release unusually high levels of an enzyme that is not adequate to maintain normal blood pressure (12). We hypothesized that such persistent stimulation of renin cells would lead to arteriolar disease -and given the co-inductive relationship between renin cells and nerve fibers - would be accompanied by arterial hyperinnervation (10). To address this hypothesis, we performed 3D imaging of *Ren1^c-tdTomato/tdTomato^* mice at 1-, 3-, and 7- months of age. As suspected, there was progressive concentric hypertrophy and sympathetic hyperinnervation that intensified over time (Figure 7A, Supplemental Figure 5 and Video 9). Renin cells labeled with tdTomato increasingly exhibited a distinct bobbin-like arrangement, forming concentric layers around SMCs over time. Hypertrophy of arterioles was particularly severe near the glomeruli, eventually encircling them entirely. Examination of sagittal and coronal images of hypertrophied AAs from 7-month-old mice revealed an additional inner layer of αSMA-positive SMCs lining the arterial lumen, further narrowing it (Figure 7B, Supplemental Figure 6). Simultaneously, substantial nerve fiber expansion was observed in parallel with vascular hypertrophy (Figure 7A). Specifically, numerous axonal branches closely associated with renin-stimulated cells extended in multiple directions around enlarged AAs, eventually enveloping the glomeruli. Coronal view further revealed nerve fibers interwoven among the concentric layers of renin cells (Figure 7B, Supplemental Video 10). Immunostaining of 2D kidney sections indicated that these hyper-innervated nerve fibers predominantly consisted of TH-positive sympathetic nerves, although CGRP-positive sensory nerves were also identified (Supplemental Figure 7A and B). Additionally, nerve fibers strongly expressed the synaptic marker synaptophysin, indicating active synapse formation between nerve fibers and the concentric layers of renin cells (Supplemental Figure 7C). Finally, quantitative analysis of mRNA expression levels of neurotrophins and their receptors revealed that the relative mRNA expression levels of *Ngf* and *Ntrk2* were significantly higher in the kidneys of 4–5-month-old *Ren1^c-tdTomato/tdTomato^* mice (mean ± SD; *Ngf*: 3.28 ± 0.88, p = 0.0075; *Ntrk2*: 31.37 ± 6.07, *P* <0.0001) compared to age-matched *Ren1^c-tdTomato/+^* control mice (Figure 7C, Supplemental Figure 8). Relative mRNA expression levels of *Ngf* and *Ntrk2* exhibited a significant positive correlation with age (*Ngf*: r = 0.60, *P* = 0.0287; *Ntrk2*: r = 0.86, *P* = 0.0002) (Figure 7D). These results are consistent with the histological progression of the disease and suggest that increased *Ngf* and *Ntrk2* expression are involved in the aberrant hyperinnervation and vascular remodeling associated with chronic renin enzymatic insufficiency.

**Figure 7.**
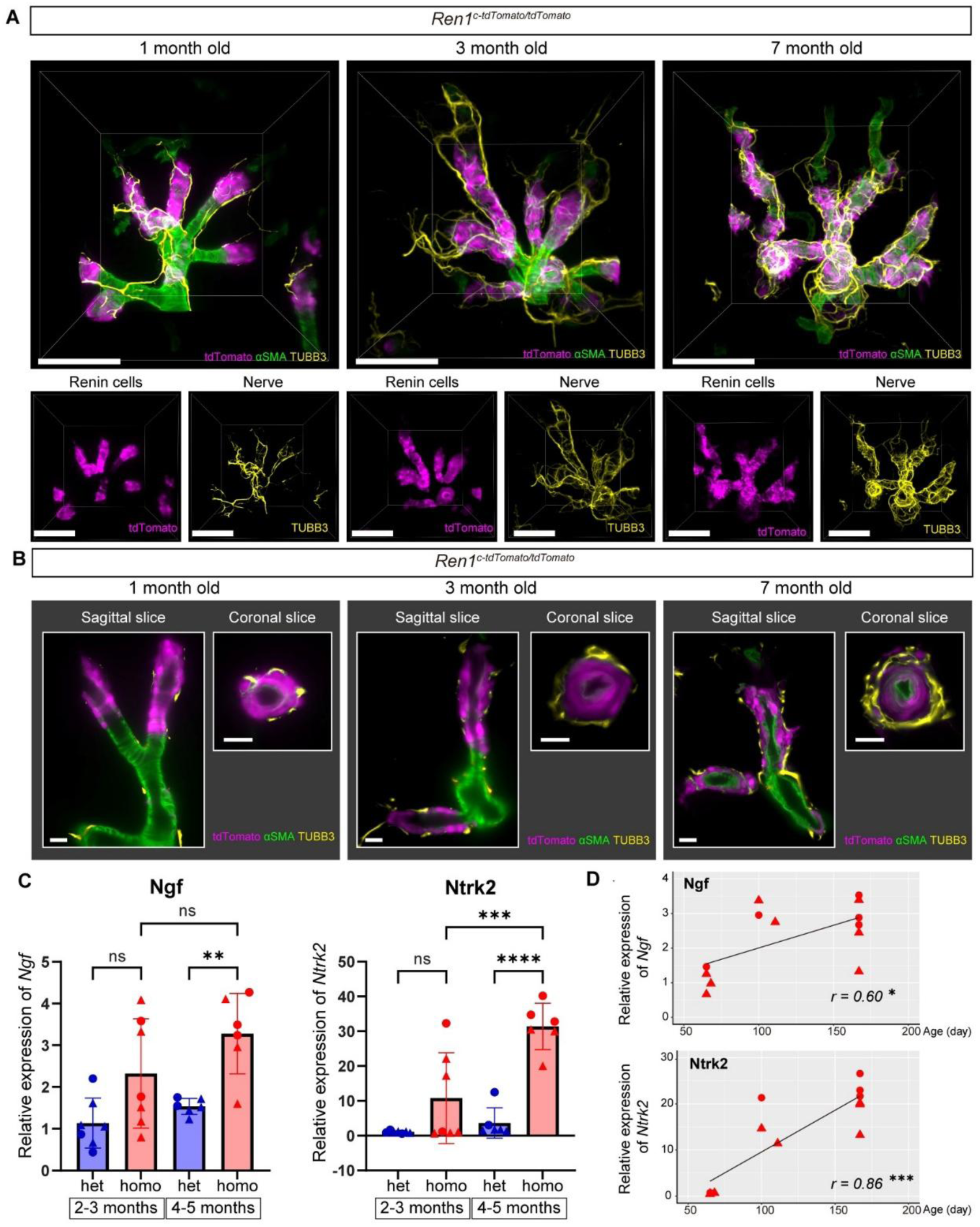
Chronic RAS inhibition drives concentric arterial hypertrophy and sympathetic hyperinnervation. (A) Representative 3D images of renin cells, vascular mural cells, and nerve fibers in kidneys of *Ren1^c-tdTomato/tdTomato^* (homo) mice at 1, 3, and 7 months of age. Renin cells exhibit concentric expansion and progressive hyperinnervation. Lower panels separately depict renin cells and nerve fibers. Scale bars: 100 µm. (B) Sagittal and coronal slices from 3D imaging illustrating progressive concentric thickening of afferent arterioles and hyperinnervation around renin cells in *Ren1^c-tdTomato/tdTomato^* (homo) mice at 1, 3, and 7 months. Coronal slices views highlight progressive narrowing of arterial lumens and increasing complexity of nerve fiber arrangement. Scale bars: 20 µm. (C) Quantitative RT-PCR for *Ngf* and *Ntrk2* in kidneys from *Ren1^c-tdTomato/+^* (het) and *Ren1^c-tdTomato/tdTomato^* (homo) mice at 2–3 and 4–5 months of age. *Ngf* and *Ntrk2* increased significantly in 4–5-month-old homo mice compared to age-matched het controls (two-way ANOVA). Data normalized to the mean values of 2–3-month-old het mice. Triangles indicate males; circles indicate females. (D) Pearson’s correlation analysis demonstrates significant positive correlations between age and relative expression levels of *Ngf* and *Ntrk2* mRNA in *Ren1^c-tdTomato/tdTomato^* (homo) mice. Data expressed relative to the mean value of 2–5-month-old het mice. Triangles indicate males; circles indicate females. *P < 0.05, **P < 0.01, ***P < 0.001, ***P < 0.0001; ns, no significant difference See also Supplemental Videos 9 and 10.

## Discussion

Here, we visualized with exquisite details the rare renin cell in its “habitat” without disrupting the complex architecture of the kidney. Further, we show the dynamic interactions of renin cells with other cells in the arterioles, glomeruli and nerve fibers during healthy development, physiological challenges and arterial disease in a novel model of renin enzymatic insufficiency (Figure 8).

**Figure 8.**
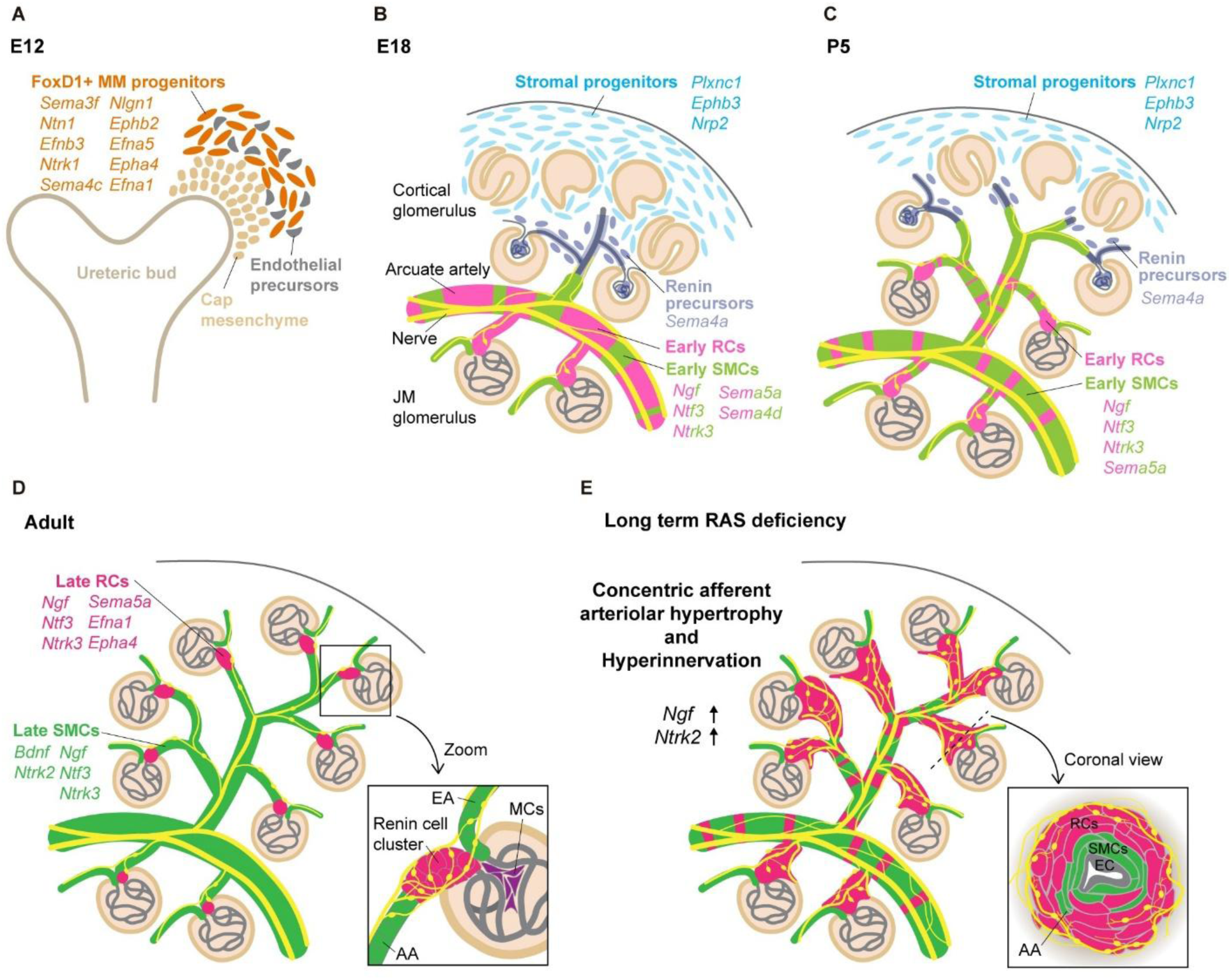
Developmental dynamics and pathological remodeling of renin cells, renal vasculature, and nerve growth during normal development and renin enzymatic deficiency. (A) At E12, FoxD1-positive metanephric mesenchymal (MM) progenitors express axon guidance molecules involved in early neurovascular patterning. (B) At E18, stromal progenitors, renin precursors, early renin cells (RCs), and early smooth muscle cells (SMCs) coordinate arterial formation and innervation. Early RCs/SMCs highly express neurotrophic factors such as *Ngf* and *Ntf3*, which stabilize and maintain tissue innervation. Arcuate arteries are extensively covered by RCs. Juxtamedullary afferent arterioles (JM-AAs) and arcuate arteries are extensively covered by RCs, with nerves extending along them. In contrast, nerve fibers extend ahead of (and prior to) the vascular wall cells in cortical arteries. (C) At P5, early RCs diminish in arcuate arteries but remain prominent in JM-AAs. In the cortex, early RC clusters emerge at cortical AA tips connected to developing glomeruli, forming striped patterns along arterial walls. In peripheral cortical regions, nerves extend before RC differentiation. (D) In adults, mature renin cells (late RCs) localize exclusively at juxtaglomerular regions, prominently in cortical glomeruli compared to JM glomeruli. Late RCs continuously express neurotrophic factors (*Ngf, Ntf3, Ntrk3, Sema5a*), and axon guidance molecules (*Efna1, Epha4*) re-emerge at this stage. Late SMCs prominently express Bdnf and Ntrk2, alongside Ngf, Ntf3, and Ntrk3. Collectively, these expression profiles suggest cooperative roles of late RCs and SMCs in maintaining neurovascular structural integrity. Nerves branch into fine terminals forming effector junctions with RCs and extend toward efferent arterioles without innervating mesangial cells (MCs). (E) Long-term renin-angiotensin system (RAS) deficiency incudes concentric afferent arteriolar hypertrophy and hyperinnervation, associated with increased expression of *Ngf and Ntrk2*, which likely contributes to structural remodeling of the renal vasculature and neural network. The coronal view illustrates concentric accumulation of SMCs, enlarged RCs encroaching upon the vascular wall, and proliferating nerve fibers, resulting significant lumen narrowing. EC, endothelial cells.

Due to their scarcity (0.01 % of the total kidney cell mass), dynamic distribution, and complex spatial interactions with glomeruli and macula densa, renin cells present unique challenges for conventional 2D histological imaging. To overcome those limitations, we utilized the *Ren1^c-tdTomato^* mouse model, which express an ultra bright fluorescent reporter, tdTomato, under the endogenous renin gene locus (12). Combining this model with tissue clearing allowed unprecedented, in depth and fine details of the renin cells morphology, localization, and cell-cell interactions throughout development and in response to physiological challenges and pathological stimuli. Our study significantly advances knowledge largely obtained through histological assessments (9, 21) and meticulous microdissections of the renal arterial tree (3). For instance, in the latter, we classified the distribution of renin cells within AAs into seven distinct types which were corroborated by the present study. However, the proportion of AAs totally lacking renin cells (type 5) was over 30% in the previous study (3). Applying such classification to our 3D data, we found that only 2.5% of AAs in adult mice lacked renin cells under normal conditions. These findings illustrate the dramatically improved detection efficiency achieved with the newer methodology.

We also found differences in the timing and distribution of renin cells located in juxtamedullary arterioles versus those present in cortical AAs. In the non-stressed normal adult animal, renin clusters -the group of JG cells located near the glomeruli-were larger in cortical AAs, indicating that these cortical regions likely serve as the primary sites of renin production. There are notable differences in pressure load and vascular tone between juxtamedullary AAs and cortical AAs (22). Juxtamedullary AAs arise from the initial portion of the cortical radiate arteries or directly from arcuate arteries and are thicker and shorter (23). Thus, juxtamedullary AAs must maintain strong vascular tension to maintain a substantial pressure gradient over the short distance between the arcuate arteries and the glomeruli (22). In contrast, and according to Bernoulli’s principle (24), the cortical circulation experiences a more gradual pressure decrease along the longer vascular path comprising the cortical radiate arteries and Aas (22). As a result, cortical AAs operate at significantly lower perfusion pressures than juxtamedullary AAs (25). Furthermore, juxtamedullary and cortical AAs have different responsiveness to Ang II: cortical AAs are more sensitive to Ang II (26, 27). Thus, it is plausible that cortical glomeruli, which must respond promptly to systemic changes such as hypotension, sodium depletion, and reductions in Ang II, require greater renin-mediated regulatory capacity, thereby maintaining a larger population of renin cells in cortical AAs. Interestingly, under short-term captopril treatment with low sodium diet discussed below, the marked difference in renin cluster size between juxtamedullary and cortical AAs disappeared, suggesting recruitment of renin cells also occurs actively within juxtamedullary AAs in response to physiological stress indicating that renin cells in juxtamedullary AAs are sensitive and respond to the systemic hypotension and fluid depletion known to occur in this experimental design (5–7). It remains to be determined whether the pressure-sensing mechanism in these cells is equally efficient to the baroreceptor of cells located in cortical vessels.

Sympathetic stimulation via β1 adrenergic receptors on renin cells is one of the primary mechanisms promoting renin production and secretion (28). However, conventional 2D imaging techniques have limited our understanding of the precise spatial and temporal interactions between renin cells and nerve fibers. Here, we uncovered dynamic developmental changes in the spatial distribution and innervation patterns of renin cells. Between E17 and P0, renin-expressing cells extensively covered cortico-medullary and arcuate arteries, as well as juxtamedullary AAs. In contrast, cortical AAs prominently expressed renin primarily within JG regions connected to mature glomeruli, whereas peripheral cortical arterioles exhibited sparse or minimal renin expression that was often below detection thresholds. Importantly, nerve fibers were observed extending along immature arterial walls derived from FoxD1-positive progenitor cells prior to detectable renin expression, suggesting that those progenitors and developing axons guide each other and actively influence renal arterial and renin cell maturation.

An initially paradoxical result from our developmental analysis was that despite cortical AAs -which have lower perfusion pressure and higher sensitivity to physiological changes-being the primary sites of renin production in adult kidneys, widespread renin expression during fetal development predominantly occurred in juxtamedullary AAs. Moreover, despite the expectation that blood flow would be less stable and lower in developing cortical arterioles, widespread renin expression in these vessels was not as prominent as in juxtamedullary AAs. We propose the following explanations for this apparent paradox. These possibilities are not contradictory but may exist simultaneously, each contributing to different aspects of renal development and functional maturation. First, renin cells participate in the morphogenesis of the vasculature which proceeds centrifugally. This may be independent of flow and systemic pressure (1, 2). Thus, the programmed centrifugal maturation -whereby juxtamedullary nephrons appear and mature earlier than cortical ones-may itself influence the developmental pattern of renin expression. In addition, it seems reasonable that in the early stages of kidney development, renin production in the thick renal arteries and the earlier-developing juxtamedullary AAs is necessary also to maintain local renal blood flow and secondarily the systemic circulation. As the animals mature and transition to extrauterine life, regulation of the systemic circulation and perfusion pressure becomes crucial to sustain extrauterine life and the more numerous and responsive cortical renin cells must join in the control of the systemic circulation. Interestingly, a study in fetal lambs showed that plasma renin levels under non-stressed conditions exceeded maternal levels and increased approximately 10-fold following acute hemorrhage (8-10% blood loss), underscoring the functional importance of the RAS in fetal circulation stability (29). Healthy full-term human infants also exhibit peak plasma renin concentrations within 1-2 days after birth, rapidly declining thereafter, likely representing RAS activation in response to the perinatal stress imposed by labor and delivery (30, 31). Similar findings were confirmed in newborn lambs, where nephrectomy abolished this transient increase, identifying the kidneys as the primary renin source (32). Furthermore, renal denervation studies in fetal lambs demonstrated critical regulation by renal nerves of perinatal renin gene expression and plasma renin (33, 34). Additionally, a previous study based on observations of 2D sections of human kidneys from stillborn fetuses showed that renin-expressing cells normally shift from the inner to outer renal cortex during gestation; however, intrauterine growth restriction disrupts this shift, maintaining renin expression in the inner cortex, likely due to altered renal blood flow and hypoxemia (35). In particular, in mouse kidneys, the timing of birth coincides with the acceleration of vascular maturation and glomerular development in the cortical region during the centrifugal maturation process. Thus, widespread fetal renin expression throughout the renal vasculature and robust renin expression along the juxtamedullary AAs may represent a programmed developmental response necessary to sustain -grow of the vasculature and- renal blood flow early in development and systemic circulation during fetal life and the perinatal circulatory transition. After birth, as systemic circulation stabilizes and the cortical glomeruli develop, renin cells localize to the JG apparatus and more precisely regulate renin production and secretion through neural input, macula densa signaling, and pressure receptor mechanisms.

The importance of innervation for renin-expressing cell differentiation, migration and arteriolar morphogenesis in development has also been suggested by previous observations from several transgenic mice models (36, 37). Sympathetic stimulation via β1 adrenergic receptors on renin cells activates the cAMP signaling pathway, critically promoting renin synthesis and secretion (38). Mice lacking the guanine nucleotide-binding protein (Gsα) specifically in renin-lineage cells show suppressed renin expression at all developmental stages and lacked renin cells throughout the renal vasculature, accompanied by reduced length and fewer branching points of AAs (36). In contrast, β1 and β2-adrenoceptor knockout mice showed a dramatic reduction in renin expression particularly within large renal vessels, however, retained normal arteriole morphology and typical JG renin localization (37). Neubauer et al. suggest that β-receptor deficiency alone does not fully recapitulate Gsα-deficient phenotypes, indicating the existence of complementary cAMP activation pathways independent of β-receptors (37). In this context, dopamine released from renal sympathetic nerve terminals is proposed as one such complementary Gsα-coupled stimulus (37). Additionally, CGRP, released from sensory nerve terminals is another neurogenic Gsα-coupled pathway capable of stimulating cAMP signaling. Indeed, our 3D imaging analyses, along with a previous elegant report (39), confirmed CGRP-positive sensory nerve fibers innervating the renal vasculature, suggesting their potential involvement in arteriolar development and renin localization.

The accurate anatomical and functional maturation of neurovascular networks require coordinated neurovascular interactions mediated by axon guidance molecules and neurotrophins (40, 41). For example, in the limb skin, peripheral nerves determine the pattern of arterial differentiation and vascular branching (42, 43). Our 3D imaging of the centrifugal maturation of renal arterioles revealed that in cortical AAs, nerve fibers extended along immature vascular walls derived from FoxD1-positive progenitor cells prior to the detection of renin expression. These observations support the possibility that nerves induced by FoxD1-derived vascular progenitor cells actively contribute to establishing the location and functional maturation of renin cells. This hypothesis is further supported by our single-cell RNA-Seq analyses, which demonstrated interactions between FoxD1-lineage vascular wall cells and nerves beyond Gsα-cAMP pathway-mediated mechanisms. Specifically, our single-cell RNA-Seq revealed that FoxD1-derived progenitors initially express axon guidance molecules for early neural guidance, later shifting to neurotrophins to stabilize mature neural connections (Figure 8). Such active participation positions FoxD1-lineage cells as crucial intermediaries that integrate developmental cues to establish neurovascular organization. At the molecular level, axon guidance molecules such as *Sema3f, Ntn1* and *Enfb3* which regulate nerve attraction and repulsion, likely play pivotal roles in orchestrating precise nerve placement and establishing proper neurovascular connections. In addition, several axon guidance signaling pathways identified in FoxD1-derived cells, including Slit/Robo, Eph/Ephrin, Semaphorins/Plexin-Neuropilins, and Netrins/UNC5, have recently been implicated in angiogenesis (44), suggesting roles in positioning and differentiation of FoxD1-derived vascular mural cells. In support, FoxD1-specific *Ntn1* knockout disrupts the patterning of renal neurovascular networks, showing endothelial cells as primary responders to Netrin-1, with nerve fibers subsequently following vascular patterns (45). Furthermore, differential expression patterns of neurotrophins and their receptors provided additional mechanistic insights. Notably, *Ngf* expression inversely correlated with its receptor *Ntrk1*, strongly suggesting that Ngf primarily sustain adjacent nerve fibers through paracrine mechanisms. Ngf is synthesized in target tissues of the sympathetic nervous system and is widely recognized as a crucial neurotrophic factor essential for the elongation, survival, and maintenance of sympathetic nerve axons (46, 47). Given that renin secretion is tightly regulated by sympathetic nerve activity, renin cells may actively support sympathetic innervation via Ngf production, establishing functional neurovascular interactions. Conversely, coordinated expression of Bdnf– Ntrk2 and Ntf3–Ntrk3 receptor-ligand pairs supports autocrine signaling or reciprocal interactions within vascular mural cells, potentially contributing to vascular remodeling and neural plasticity. Notably, *Bdnf* and *Ntrk2* expression was highly enriched in mature SMCs and pericytes but was nearly absent in mature JG cells, suggesting specific role for the Bdnf–Ntrk2 pathway in establishing and maintaining vascular mural cell identity. Understanding the biological significance of these reciprocal interactions between neural growth/innervation and FoxD1-lineage cell differentiation will provide deeper insights of neurovascular development and its implications in renal diseases.

We have shown that renin cell descendants retain the memory and plasticity of the renin phenotype (2, 48). Thus, when an adult animal is exposed to a homeostatic threat, SMCs, MCs and interstitial pericytes reenact such memory and are transformed into renin-producing cells. Here, we show the first high-resolution 3D evidence of this phenomenon. During a short-term physiological challenge such as sodium depletion and captopril treatment, the recruitment of renin cells is characterized by an increased in volume (and number) of individual cells leading to hypertrophy of each JG cell cluster. This is also accompanied by the transformation of SMCs along the arterioles into renin-producing cells. The underlying mechanism involves chromatin remodeling and rapid epigenetic changes (1, 2, 49), without cell proliferation or migration (50). Interestingly, each renin cell increases its volume, likely reflecting enhanced synthetic activity and expanded organelle compartments, including renin granules, ER, Golgi, and mitochondria—typical markers of increased renin synthesis and release (1, 2, 21). Changes in ionic and water flow may also contribute but remain unexplored.

Our study also highlights the progression of concentric afferent arteriolar hypertrophy and hyperinnervation in a novel model of renin enzymatic insufficiency using *Ren1^c-tdTomato/tdTomato^* mice, which uniquely exhibit insufficient activity of the renin enzyme (12). While prolonged RAS blockade is commonly employed to manage hypertension and chronic kidney disease, paradoxical adverse outcomes such as vascular remodeling and sympathetic hyperinnervation occur (8–10). In our recent report, we performed 3D imaging of the renal cortex in 3-month-old renin knockout mice (*Ren1^c-/-^; Ren1^c-Cre^; R26R^mTmG^*) and *SMMHC^CreERT2^; R26R^tdTomato^; Ren1^c-YFP^* mice subjected to long-term RAS inhibition through oral administration of the ACE inhibitor captopril for 6 months, demonstrating significant hyperinnervation as well as concentric afferent arteriolar hypertrophy (10). However, in *Ren1^c-Cre^* mice, all renin-lineage cells were permanently labeled irrespective of their active renin expression status, and in *Ren1^c-YFP^* mice, YFP fluorescence faded rapidly, precluding reliable 3D immunostaining. Consequently, it was difficult to simultaneously visualize the 3D structure of actively renin-producing cells, blood vessels, and nerves. In this study, we circumvented these technical limitations by utilizing newly developed *Ren1^c-tdTomato/tdTomato^* mice that express bright and stable tdTomato fluorescence in active cells that synthesize a hypoactive renin enzyme (12). We observed phenotypes similar to those induced by other forms of RAS inhibition, including vascular hypertrophy and hyperinnervation, reinforcing the hypothesis that chronic stimulation of renin cells due to prolonged RAS inhibition itself is fundamentally responsible for driving these pathological changes. Previous findings showing the absence of vascular hypertrophy in conditional renin cell ablation (51) or during chronic administration of antihypertensive medications other than RAS inhibitors (52) further strengthen this conclusion. In our previous study, we demonstrated elevated expression of *Ngf* mRNA in the JG region and renal cortical interstitial vasculature of renin knockout mice (10). Based on these findings, we hypothesized the existence of a positive feedback loop, wherein renin cells, inflammatory cells, and myofibroblasts enhance hyperinnervation through Ngf secretion, while the resulting hyperactivated sympathetic nerves further stimulate (captopril treatment) or attempt to stimulate (renin knock out) renin production (10). The significant increase in *Ngf* expression observed in the *Ren1^c-tdTomato/tdTomato^* mice in the present study provides additional evidence supporting Ngf’s role in driving hyperinnervation. Importantly, the density of sympathetic innervation typically correlates with the level of Ngf production in the target organ (53, 54). Thus, the elevated Ngf levels observed in the kidney likely play a direct and critical role in mediating the pathological sympathetic hyperinnervation identified in this study. In addition, our study uncovered significant upregulation of *Ntrk2* expression in hypertrophied renal vasculature, suggesting a critical involvement of the Bdnf–Ntrk2 pathway in vascular remodeling. Given that Ntrk2 signaling similarly regulates vascular SMC proliferation and extracellular matrix production, enhanced *Ntrk2* expression likely acts synergistically with Ngf to further potentiate the arterial hypertrophy, the embryonic matrix-secretory phenotype (9), and the hyperinnervation observed in our model. The concerted elevation of Ngf and Ntrk2 thus identifies these pathways as essential mediators of the observed pathological changes. Further studies using genetic modifications and pharmacological inhibition will clarify the role of Ngf and Ntrk2 signaling in pathological remodeling as well as renal neurovascular development.

In conclusion, our study provides a high-resolution map of the dynamic spatial relationships between renin cells, renal vasculature, and sympathetic innervation across developmental and disease states. As renin cells are integral to morphogenesis of the kidney vasculature, blood pressure regulation and fluid homeostasis, elucidating their ontogeny, regulatory cues, and pathological remodeling has both fundamental and clinical significance. These findings advance our understanding of renovascular biology and offer potential avenues for the development of antihypertensive strategies and kidney regeneration and bioengineering, including organoid modeling from induced pluripotent stem cells.

## Methods

Further information can be found in Supplemental Methods.

### Sex as a biological variable

Our study examined male and female animals, and similar findings are reported for both sexes.

### Animals

To label renin-expressing cells with fluorescent tdTomato under the control of the endogenous renin gene environment, we used *Ren1^c-T2A-tdTomato^*(*Ren1^c-tdTomato^*) mice generated in our laboratory (12). Heterozygous mice are indicated as *Ren1^c-tdTomato/+^*, and homozygous mice are indicated as *Ren1^c-tdTomato/tdTomato^*. *Ren1^c+/−^;Ren1^c-Cre^;R26R^mTmG^* mice were generated as previously reported (8). In this line, renin lineage cells express GFP following Cre-mediated recombination, while non-recombined cells retain RFP expression. To generate *FoxD1^GC^; R26R^tdTomato^; Ren1^c-YFP^* mice, we used the *FoxD1^GC^*mice (Jackson Laboratory: #012463), *R26R^tdTomato^* mice (Jackson Laboratory: # 007909), and *Ren1^c-YFP^* mice (55). All animals used in this study were housed in a temperature- and humidity-controlled room under a 12-hour light/12-hour dark cycle.

### Short-term captopril treatment with low salt diet

We treated 3-month -old *Ren1^c-tdTomato/+^* (Het) mice *ad libitum* with captopril added to the drinking water (0.5 g/L) and a low sodium (0.1% Na^+^) diet for 8 days.

### Processing of mouse kidneys

For frozen sections, kidney tissues from *Ren1^c-tdTomato^*and *FoxD1^GC^; R26R^tdTomato^; Ren1^c-YFP^* mice were fixed in 4% paraformaldehyde (PFA) for 2 hours at 4°C. After washing, tissues were cryoprotected in 30% sucrose suspended in PBS overnight at 4°C, embedded in optimal cutting temperature compound (O.C.T.; Thermo Fisher Scientific), and subsequently frozen at -80°C.

### Immunofluorescence staining

Immunofluorescence staining was performed on 8 µm cryo-fixed sections. After blocking with 3% donkey serum, sections were incubated overnight at 4°C with primary antibodies. Following washing and re-blocking with 3% donkey serum, the sections were incubated for 2 hours at room temperature with secondary antibodies. After further washes, sections were treated with an autofluorescence quenching kit (Vector Laboratories), counterstained with Hoechst 33342 (Thermo Fisher Scientific) for nuclear visualization, and mounted with mounting medium (Vector Laboratories). The primary antibodies used were as follows: anti-Renin antibody (1:5000 dilution; ab212197, Abcam), anti-Renin antibody produced by Inagami et al (56). (1:5000 dilution), Alexa Fluor 647-conjugated anti-TUBB3 antibody (1:200 dilution; 657406, BioLegend, San Diego, CA), anti-TH antibody (1:200 dilution; AB152), anti-synaptophysin antibody (1:100 dilution; ab14692, Abcam), Alexa Fluor 488-conjugated anti-CGRP-I+CGRP-II antibody (1:100 dilution; ab305115, Abcam), fluorescein isothiocyanate (FITC)-conjugated anti-αSMA antibody (1:100 dilution; F3777, Sigma-Aldrich), and anti-PECAM-1 antibody (1:100 dilution; AF3628, Bio-Techne). The secondary antibodies used were Alexa Fluor 488-conjugated donkey anti-rabbit antibody (1:400 dilution; Thermo Fisher Scientific), Alexa Fluor 488-conjugated donkey anti-goat antibody (1:400 dilution; Thermo Fisher Scientific), and Alexa Fluor 647-conjugated donkey anti-rabbit antibody (1:400 dilution; Thermo Fisher Scientific).

### Microscopy

Tissue sections were imaged using a Zeiss Imager M2 microscope equipped with an Apotome.2 optical sectioning device fitted with AxioCam 305 color and AxioCam 506 mono cameras (Zeiss).

### Tissue clearing and whole-mount immunostaining of mouse kidneys

Mouse kidney tissue clearing and whole-mount immunostaining were conducted using an optimized version of the Clear, Unobstructed Brain/Body Imaging Cocktails and Computational Analysis (CUBIC) method (57). Procedures were based on previously described protocols (13, 58). Mice were anesthetized using tribromoethanol (300 mg/kg) and subsequently perfused through the left ventricle with 20 ml PBS, followed by 30 ml of 4% PFA. Kidneys were excised, bisected, and fixed overnight in 4% PFA. All subsequent procedures were conducted with gentle agitation. After fixation, samples were washed in PBS for 6 hours, followed by immersion in CUBIC-L solution [10 wt% N-butyldiethanolamine and 10 wt% Triton X-100 (MilliporeSigma)] at 37°C for 5 days.

Samples underwent another 6-hour PBS wash and were blocked overnight at room temperature (RT) in blocking buffer (PBS containing 1.0% bovine serum albumin and 0.01% sodium azide). Next, samples were incubated in immunostaining buffer (PBS containing 0.5% Triton X-100, 0.25% bovine serum albumin, and 0.01% sodium azide) supplemented with fluorescently-labeled antibodies: FITC conjugated anti-αSMA antibody (1:100 dilution, F3777, Sigma-Aldrich), Alexa fluor 647-conjugated anti-TUBB3 antibody (1:200 dilution, 657406, BioLegend), Alexa Fluor 488-conjugated anti-CGRP-I+CGRP-II antibody (1:100 dilution, ab305115, Abcam) or TOPRO3 nuclear dye (1:1000 dilution, T3005, Life Technologies), for 5-14 days at RT. Staining duration varied depending on sample size. After a subsequent 6-hour PBS wash, only samples stained with anti-TUBB3 antibody underwent post-fixation in 1% PFA for 3 hours. All samples were then immersed overnight at RT in a 1:1 dilution of CUBIC-R+ solution [T3741 (containing 45 wt% 2,3-dimethyl-1-phenyl-5-pryrazolone, 30 wt% nicotinamide, and 5 wt% N-butyldiethanolamine), Tokyo Chemical Industry], followed by undiluted CUBIC-R+ solution immersion at RT for an additional 2 days.

### Light-sheet fluorescence microscopy

Cleared kidney samples were imaged using a Zeiss Lightsheet 7 fluorescence microscope (Zeiss). Specimens were mounted in a refractive index matching solution (RI = 1.520; M3294, Tokyo Chemical Industry) and visualized using Lightsheet 7 detection optics with either a 5× / 0.16 foc or a Clr Plan-Neofluar 20× / 1.0 Corr nd = 1.53 objective lens. Voxel resolution was as follows: for the 5× objective lens (zoom: 1.5), x = 0.632 µm, y = 0.632 µm, and z = 3.38–3.61 µm; for the 20× objective lens (zoom range: 0.36–2.0), x = 0.118–0.656 µm, y = 0.118–0.656 µm, and z = 0.549–1.00 µm. Excitation wavelengths were 488 nm for GFP, FITC, Alexa Fluor 488, or YFP; 561 nm for tdTomato; and 638 nm for Alexa Fluor 647 or TOPRO3 signals.

### Three-dimensional image

Three-dimensional reconstruction and analysis of captured images were performed using Imaris software (Version 10.0.0, Bitplane). Raw Zeiss microscopy data (.czi files) were converted to Imaris-compatible files (.ims format) using Imaris File Converter 10.0.0, with tile scan images stitched by Imaris Stitcher 10.0.0. Image processing in Imaris was performed according to established protocols (13, 58). Three-dimensional reconstructions were cropped into areas of interest using the crop function, and images or videos were generated using the snapshot and animation functionalities, respectively.

### Dot plot for single-cell RNA-Seq analysis

Our previously published single-cell RNA-Seq data from embryonic and adult *FoxD1^Cre^; R26R^mTmG^* mouse kidneys (GSE218570 (4, 18)) was reprocessed for dot plot visualization of neurogenesis related genes in renin-lineage cells. We extracted the raw gene count matrices from each dataset and merged them using the intersection of gene symbols (59). The raw expression matrix was log-normalized and z-score transformed, and the percentage of cells expressing each neurogenesis related gene was calculated across all cell populations in the gene expression matrix that composed the renin cell trajectory. The output from this function was then plotted using ggplot2 (60).

### Pathway enrichment analysis

To calculate single-cell enrichment of gene sets and pathways, we utilized annotated gene sets from the Molecular Signatures Database (61, 62) for mouse hallmark, regulatory target, curated, cell signature, biological process, cellular component, and molecular function gene sets. Gene sets were scored on individual cells using the Single Pathway analysis in Single Cells (SiPSiC v.1.4.3) R package (19). Individual cell pathway scores for each cell population were plotted using ggplot2 (60). Significant differences were calculated by pairwise t-tests with the Benjamini-Hochberg multiple testing procedure.

### Statistics

Statistical analyses were performed using GraphPad Prism version 10.4.1 (GraphPad Software). The normality of data for unpaired comparisons was assessed by the Shapiro–Wilk test. Data confirmed as normally distributed were analyzed using Student’s t-tests, whereas non-normally distributed data were analyzed using the Mann–Whitney U test. Data are presented as mean ± standard deviation (SD) if normally distributed, or as median with interquartile range (IQR) if non-normally distributed. For multiple comparisons, pairwise t-tests with Benjamini–Hochberg correction or two-way ANOVA were used. Pearson’s correlation coefficient was calculated for linear correlation analyses. P-values <0.05 were considered statistically significant. Schematics were created with BioRender.com.

### Study approval

All procedures were performed in accordance with the National Institutes of Health guidelines for the care and use of laboratory animals and were approved by The Animal Care and Use Committee of the University of Virginia.

### Data availability

The single-cell RNA sequence data have been uploaded to the Gene Expression Omnibus (GEO) with the accession IDs GEO: GSE218570. Any additional data that support the findings of this study are available from the corresponding author upon reasonable request.

## Author contributions

MLSSL and RAG designed the study and supervised the project; MY and HY performed experiments; MY, HY and KT developed methodology; SM developed mouse model; MY, HY, JPS and SH analyzed the data; SMW performed EM; MY drafted the initial version of the manuscript; AGM, MLSSL, and RAG read, reviewed, redrafted, and edited the manuscript; HY, JPS, LFA, DM, AGM, SMW, SH, KT, SM, MLSSL and RAG reviewed and edited the manuscript; and all authors approved Its final version.

## Acknowledgments

We thank Thomas Wagamon, Fang Xu, Ahana Sinharoy, Jolie Le and Drishti Ashok Daga for excellent technical assistance. This work used ZEISS Lightsheet7 and Imaris software in the Advanced Microscopy Facility, supported by the University of Virginia School of Medicine, Research Resource Identifiers (RRID): SCR_018736. This work used F20 and sample preparation for electron microscope imaging in the Molecular Electron Microscopy Core, which is supported by the University of Virginia School of Medicine, Research Resource Identifiers (RRID):SCR_019031, which is supported in part by the School of Medicine and built with NIH grant G20-RR31199.

Foundation; supported by National Institutes of Health grants P50 DK 096373 and R01 DK 116718 to RAG, P50 DK 096373 and R01 HL148044 to MLSSL and the Japan Society for the Promotion of Science Overseas Research Fellowships to MY.

## Notes

**Conflict of interest** KT is a co-inventor on Japanese and international patents and patent applications covering CUBIC-related tissue-clearing technology. Other authors declare no competing interests.

### Competing Interest Statement

KT is a co-inventor on Japanese and international patents and patent applications covering CUBIC-related tissue-clearing technology.
Other authors declare no competing interests.

