## Supplemental materials for "Renin Cells Drive Kidney Neurovascular Development and Arterial Remodeling when Renin Activity is Deficient"

This material includes the following:

Supplemental Figures 1-8

Supplemental Video legends 1-10

Supplemental Methods

Supplemental Reference

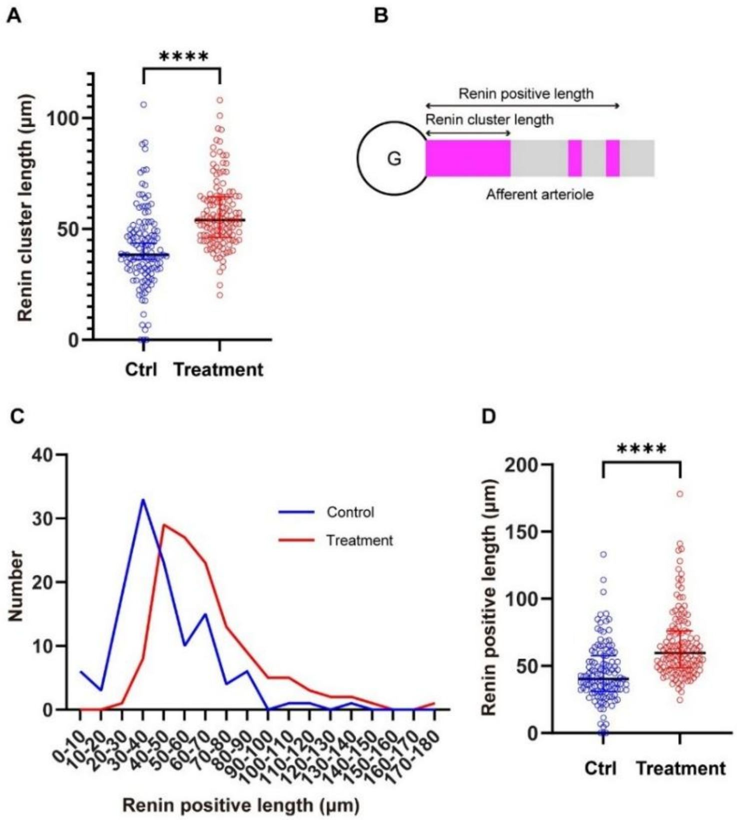


**Supplemental Figure 1. Quantitative analysis of renin cluster length and renin-positive length.**

(A) Comparison of renin cluster lengths between control (Ctrl; median 38.4 µm, IQR 31.3–49.5 µm) and treatment groups (median 53.9 µm, IQR 46.3–64.6 µm). Data points represent individual measurements from two mice per group. Statistical analysis by Mann–Whitney U test (****P < 0.0001).

(B) Schematic illustration of an afferent arteriole showing the definitions of "renin cluster length" and "renin-positive length." Renin cluster length is defined as the continuous length of the renin-expressing region extending proximally from the glomerulus (G). Renin-positive length includes the total extent of renin expression along the arteriole, encompassing discontinuous or striped patterns beyond the continuous cluster region.

(C) Histogram comparing the distribution of renin-positive lengths between control and treatment groups.

(D) Comparison of renin-positive lengths between control (median 40.3 µm, IQR 31.2–57.6 µm) and treatment groups (median 59.6 µm, IQR 48.4–76.1 µm). Data represent measurements from two mice per group. Statistical analysis by Mann–Whitney U test (****P < 0.0001).

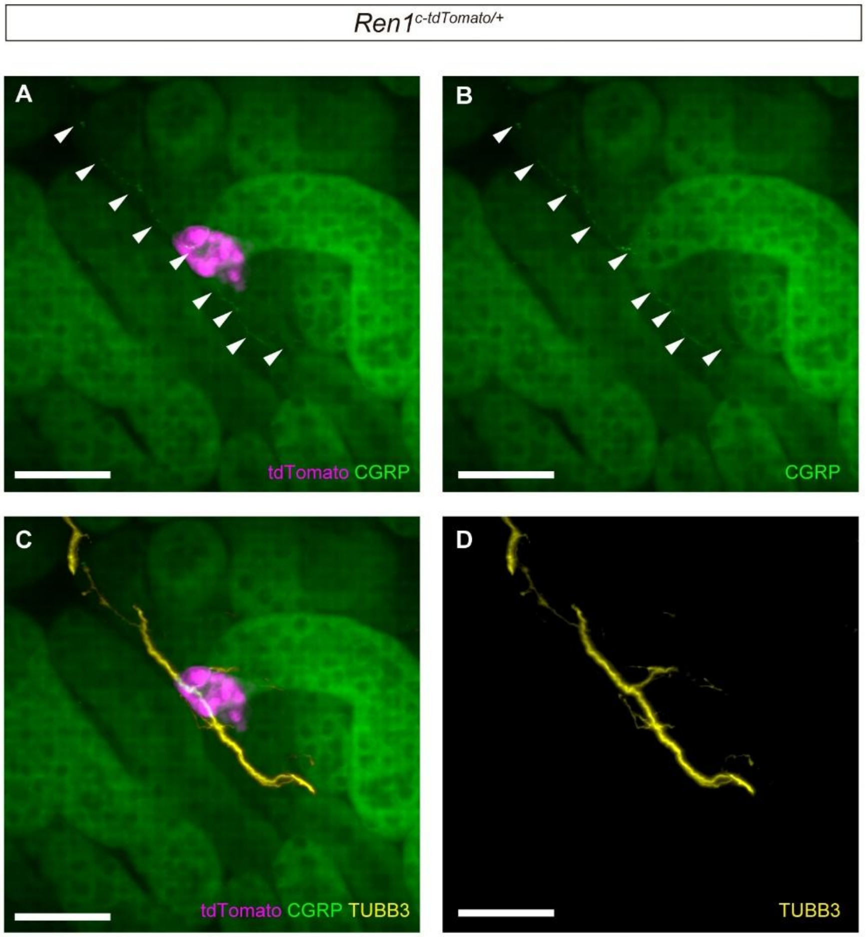


**Supplemental Figure 2. Three-dimensional images showing sensory innervation in the renal cortex of adult *Ren1$^{c\text{-}tdTomato/+}$* mice.**

(A) Immunostaining for calcitonin gene-related peptide (CGRP, sensory neuron marker, green) reveals thin sensory nerve fibers (arrowheads) closely associated with renin cells (tdTomato, magenta).

(B) CGRP staining alone is shown to highlight sensory nerve fibers. Note that due to weak fluorescence intensity of Alexa Fluor 488-conjugated CGRP antibody, background autofluorescence from renal tubules is also detected.

(C, D) Double-labeling of CGRP (green) and TUBB3 (pan-neuronal marker, yellow) confirms that CGRP-positive sensory fibers co-localize with TUBB3-positive nerve fibers associated with renin cells.

All images are XY-plane view (30 μm thickness). Scale bars: 50 μm.

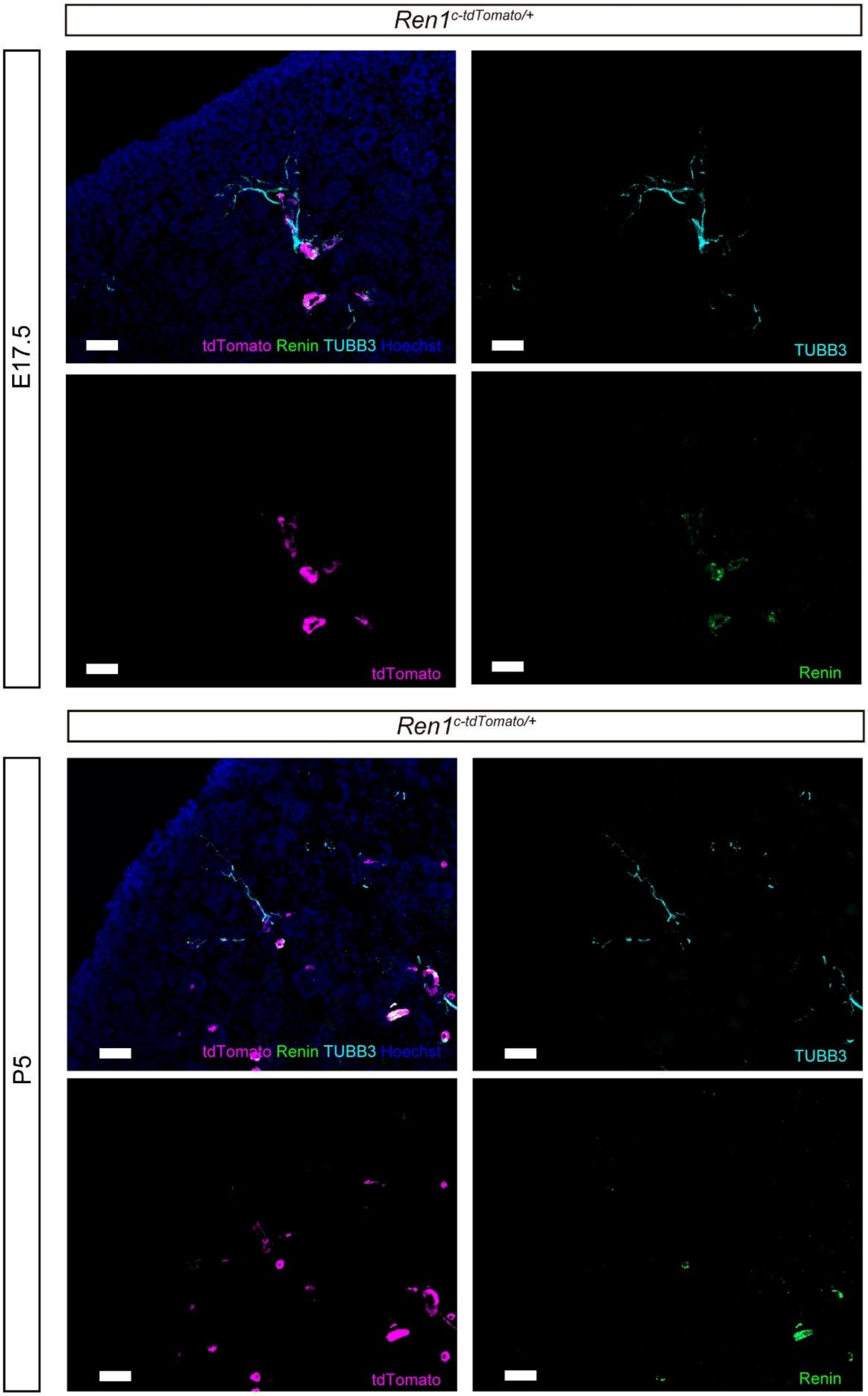

*Ren1*$^{c\text{-}tdTomato/+}$
E17.5
tdTomato Renin TUBB3 Hoechst
TUBB3
tdTomato
Renin
*Ren1*$^{c\text{-}tdTomato/+}$
P5
tdTomato Renin TUBB3 Hoechst
TUBB3
tdTomato
Renin

**Supplemental Figure 3. Nerve fibers extend beyond renin-expressing cells during renal arteriole development.**

Immunofluorescence staining of kidney sections from E17.5 (A) and P5 (B) *Ren1*$^{c\text{-}tdTomato/+}$ mice. Sections were stained for renin protein (green) and neuronal marker TUBB3 (cyan). Nuclei were counterstained with Hoechst (blue). Renin protein signals co-localized closely with tdTomato (magenta) reporter expression. At both developmental stages, nerve fibers (TUBB3-positive) extended further distally beyond the renin-positive vascular wall cells.

Scale bars: 50 μm.

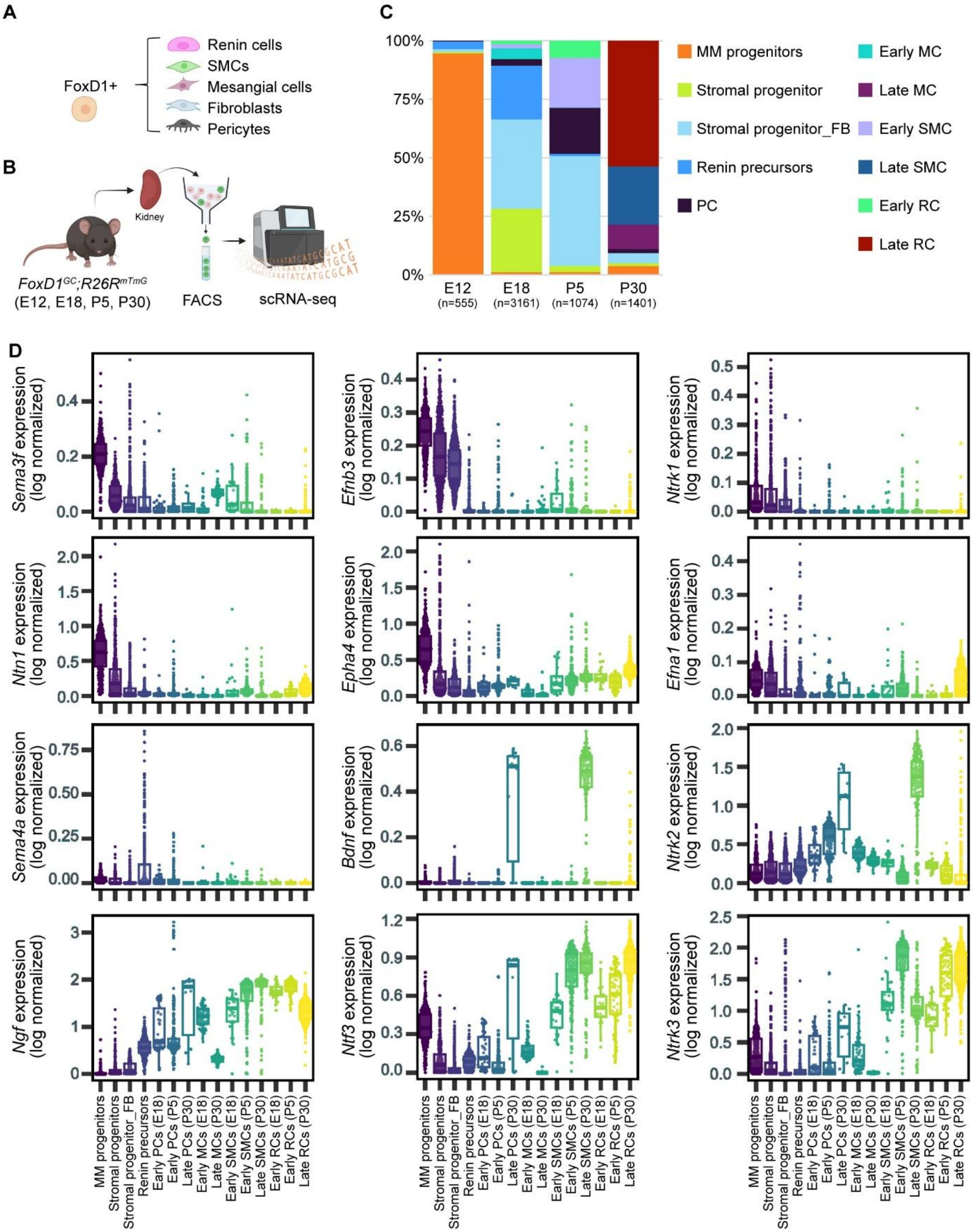
A
FoxD1+
Renin cells
SMCs
Mesangial cells
Fibroblasts
Pericytes
B
Kidney
FoxD1^GC;R26R^mTmG
(E12, E18, P5, P30)
FACS
scRNA-seq
C
100%
75%
50%
25%
0%
E12 (n=555)
E18 (n=3161)
P5 (n=1074)
P30 (n=1401)
MM progenitors
Stromal progenitor
Stromal progenitor_FB
Renin precursors
PC
Early MC
Late MC
Early SMC
Late SMC
Early RC
Late RC
D
Sema3f expression (log normalized)
Efnb3 expression (log normalized)
Ntrk1 expression (log normalized)
Ntn1 expression (log normalized)
Epha4 expression (log normalized)
Efna1 expression (log normalized)
Sema4a expression (log normalized)
Bdnf expression (log normalized)
Ntrk2 expression (log normalized)
Ngf expression (log normalized)
Ntf3 expression (log normalized)
Ntrk3 expression (log normalized)
MM progenitors
Stromal progenitors
Stromal progenitor_FB
Renin precursors
Early PCs (E18)
Early PCs (P5)
Late PCs (P30)
Early MCs (E18)
Late MCs (P30)
Early SMCs (E18)
Early SMCs (P5)
Late SMCs (P30)
Early RCs (E18)
Early RCs (P5)
Late RCs (P30)

**Supplemental Figure 4. Single-cell RNA sequencing of *FoxD1$^{GC}$; R26R$^{mTmG}$* mouse kidneys across developmental stages.**

(A) FoxD1-positive stromal progenitors differentiate into renin cells, smooth muscle cells (SMCs), mesangial cells, fibroblasts, and pericytes.

(B) Schematic illustration of the experimental workflow for single-cell RNA sequencing. Kidneys from *FoxD1$^{GC}$; R26R$^{mTmG}$* mice at embryonic day 12 (E12), embryonic day 18 (E18), postnatal day 5 (P5), and postnatal day 30 (P30) were harvested. GFP-positive cells, representing FoxD1-lineage cells, were isolated via fluorescence-activated cell sorting (FACS), and single-cell RNA sequencing was subsequently performed.

(C) Alluvial plot showing the proportional distribution of FoxD1-lineage-derived cell types at each developmental stage. The numbers in parentheses represent the total number of cells analyzed at each time point.

(D) Violin plots showing stage-specific and cell type-specific expression profiles of selected nerve-related genes within FoxD1-lineage cells across developmental stages. The expression levels of axon guidance molecules (*Sema3f, Efnb3, Ntn1, Epha4, Sema4a, Efna1*), neurotrophins (*Ngf, Bdnf, Ntf3*), and their corresponding receptors (*Ntrk1, Ntrk2, Ntrk3*) are displayed. Expression values represent log-normalized counts. Each dot corresponds to a single cell, and the shape of each violin reflects the distribution of gene expression levels within each identified cell population. Cell populations are annotated at the bottom, and developmental stages are indicated within parentheses.

MM progenitors, metanephric mesenchyme progenitors; FB, fibroblast; MC, mesangial cell; RC, renin cell; PC, pericyte.

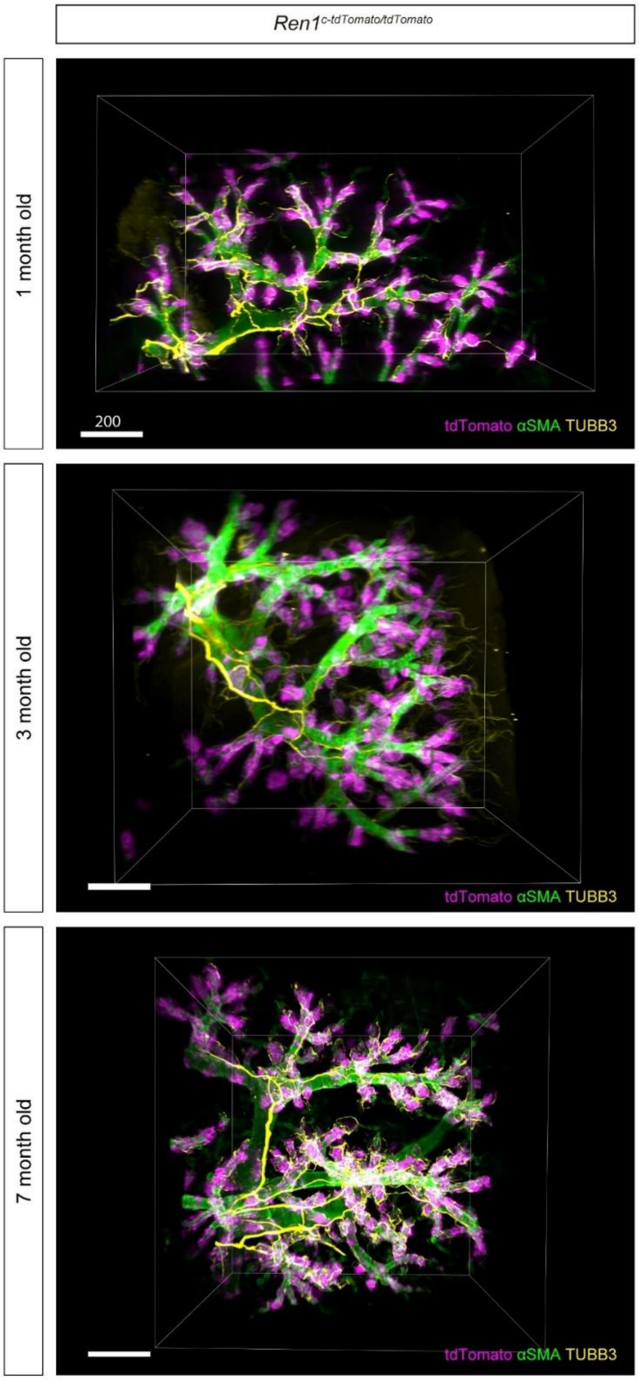


**Supplemental Figure 5. Diffuse and progressive concentric vascular hypertrophy accompanied by marked hyperinnervation is observed in the vascular tree under chronic RAS deficiency.**

Representative low-magnification 3D images illustrating progressive morphological changes in renin cells (tdTomato, magenta), vascular mural cells (αSMA, green), and nerve fibers (TUBB3, yellow) at 1, 3, and 7 months of *Ren1*$^{c\text{-}tdTomato/tdTomato}$ mice. Scale bars: 200 μm.

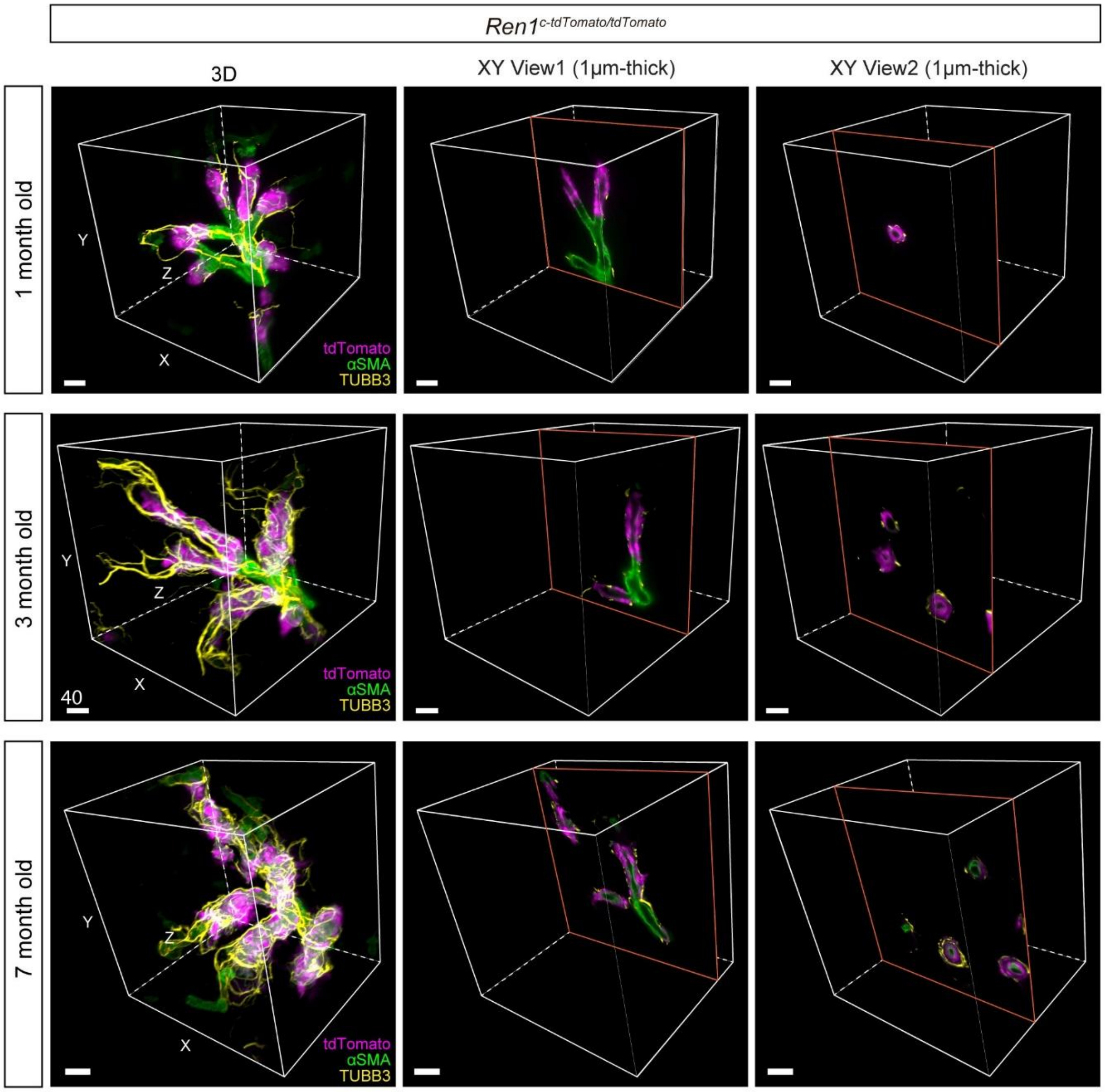


**Supplemental Figure 6. Cutting planes for XY-view sections derived from 3D reconstructions of renal afferent arterioles from *Ren1*$^{c\text{-}tdTomato/tdTomato}$ mice at 1, 3, and 7 months of age.**

Representative 3D reconstructed images demonstrate the spatial arrangement of renin cells (tdTomato, magenta), smooth muscle cells (αSMA, green), and nerve fibers (TUBB3, yellow). The middle panels illustrate the precise positions of 1 μm-thick virtual sagittal sections of afferent arterioles, while the right panels depict the positions of corresponding 1 μm-thick virtual coronal sections.

Scale bars: 40 μm.

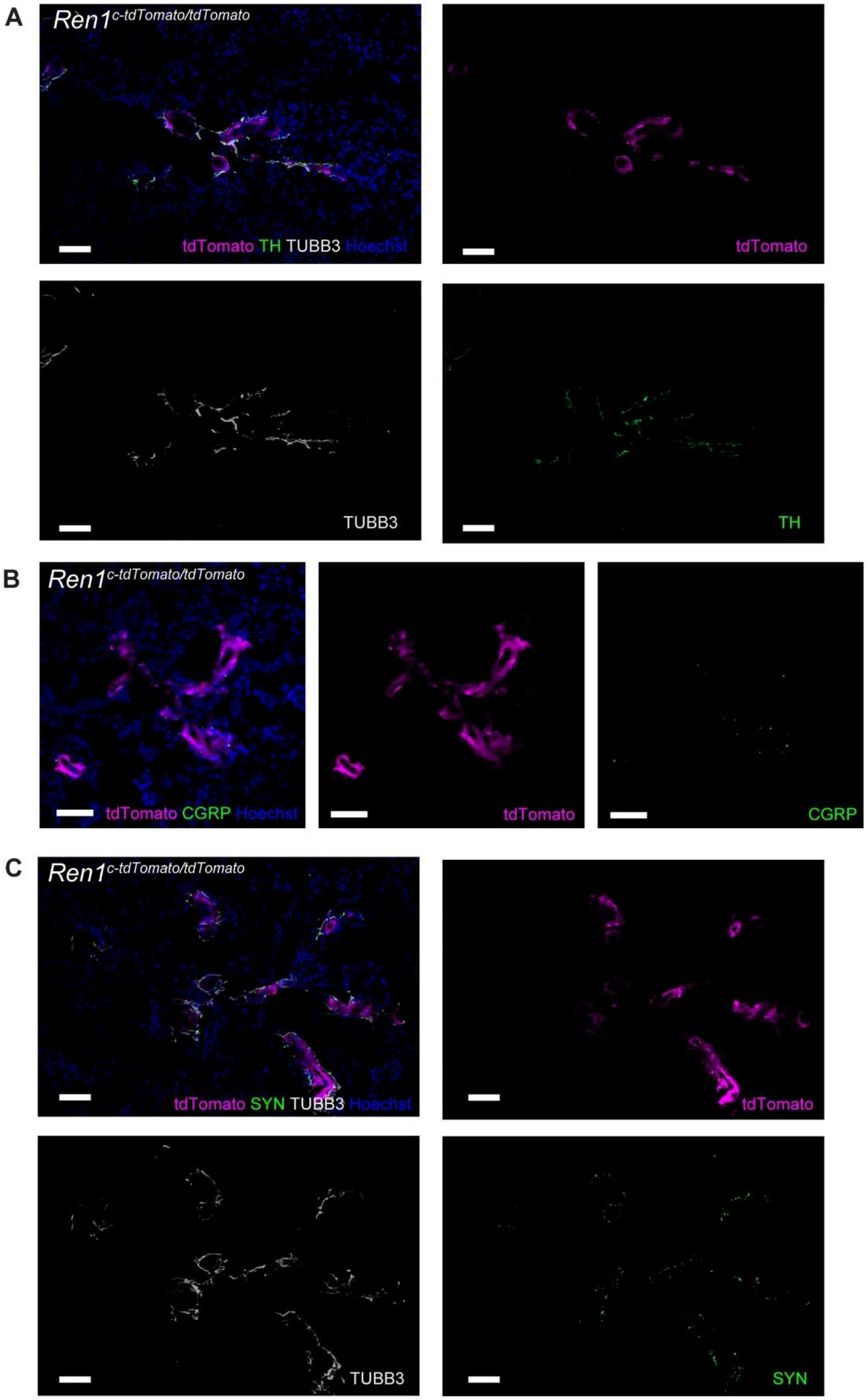
A
Ren1c-tdTomato/tdTomato
tdTomato TH TUBB3 Hoechst
tdTomato
TUBB3
TH
B
Ren1c-tdTomato/tdTomato
tdTomato CGRP Hoechst
tdTomato
CGRP
C
Ren1c-tdTomato/tdTomato
tdTomato SYN TUBB3 Hoechst
tdTomato
TUBB3
SYN

**Supplemental Figure 7. Hyperinnervation consists mainly of sympathetic nerves.**

Immunofluorescence staining of kidney sections from *Ren1*$^{c\text{-}tdTomato/tdTomato}$ mice.

(A) Immunofluorescence staining using antibodies against tyrosine hydroxylase (TH, green) and TUBB3 (white) confirmed excessive sympathetic innervation (TH-positive fibers) of renin cells (tdTomato, magenta) associated with concentric vascular hypertrophy. Nuclei were stained with Hoechst (blue).

(B) Immunofluorescence staining for calcitonin gene-related peptide (CGRP, green), a sensory nerve marker, revealed thin sensory nerve fibers closely associated with renin cells (tdTomato, magenta). Nuclei were stained with Hoechst (blue).

(C) Immunofluorescence staining with synaptophysin (SYN, green), a synaptic marker, demonstrated co-localization with TUBB3-positive nerve fibers (white), suggesting functional synaptic connections with renin cells (tdTomato, magenta) forming vascular hypertrophy. Nuclei were stained with Hoechst (blue).

Scale bars: 50 µm.

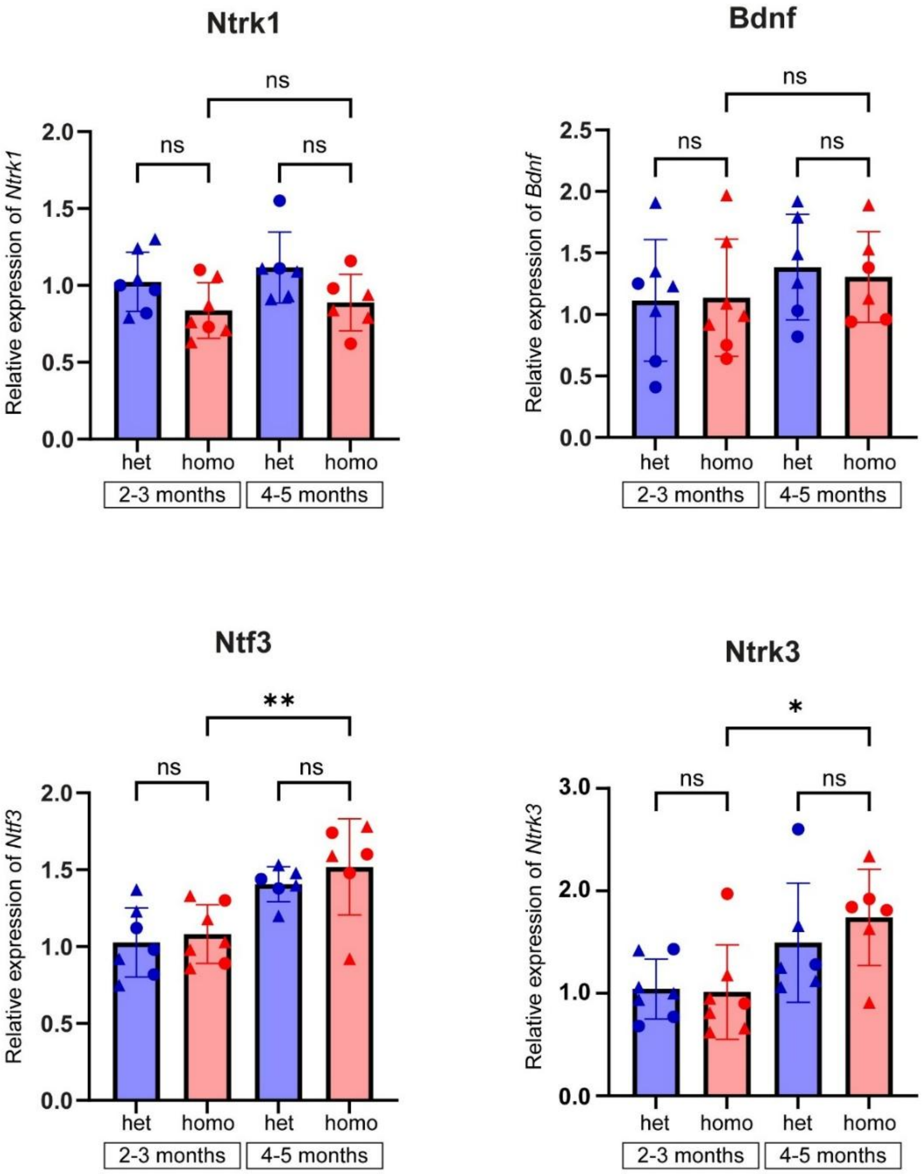


**Supplemental Figure 8. Quantitative RT-PCR analysis of *Ntrk1, Bdnf, Ntf3*, and *Ntrk3* mRNA expression in renal cortex from *Ren1*$^{c\text{-}tdTomato/+}$ (het) and *Ren1*$^{c\text{-}tdTomato/tdTomato}$ (homo) mice at 2–3 and 4–5 months of age.**

No significant differences were observed in the expression of *Ntrk1* and *Bdnf* between homo mice and age-matched het controls (two-way ANOVA). Although significant age-dependent increases in the expression of *Ntf3* and *Ntrk3* were detected in homo mice, no significant differences were observed between homo mice and age-matched het control mice (two-way ANOVA). Data are presented as relative expression normalized to the mean values of 2–3-month-old het mice. Triangles indicate males; circles indicate females.

*P < 0.05, **P < 0.01; ns, not significant.

**Supplemental Video 1. Renin cells cluster at tips of renal afferent arterioles, related to Figure 1.**

Three-dimensional structure of renin cells (tdTomato, magenta) and renal arterial tree (αSMA, green) in the kidney cortex of a $Ren1^{c\text{-}tdTomato/+}$ mouse.

**Supplemental Video 2. Cellular arrangement in afferent arterioles, related to Figure 1.**

Continuous cross-sectional views of spatial relationships among renin cells (tdTomato, magenta), smooth muscle cells (αSMA, green), and nuclei (TOPRO3, white) in the kidney cortex of a $Ren1^{c\text{-}tdTomato/+}$ mouse.

**Supplemental Video 3. Sympathetic nerves associate intimately with renin cell clusters, related to Figure 3.**

Three-dimensional visualization of nerve fibers (TUBB3, yellow), renin cells (tdTomato, magenta), and arterial structures (αSMA, green) in the kidney cortex of a $Ren1^{c\text{-}tdTomato/+}$ mouse.

**Supplemental Video 4. Sympathetic nerves branch along afferent arterioles without entering glomeruli, related to Figure 3.**

Continuous cross-sectional visualization of renin-lineage cells (GFP, green) in renal arterioles and nerves (TUBB3, cyan) of a $Ren1^{c+/-}$; $Ren1^{c\text{-}Cre}$; $R26R^{mTmG}$ mouse.

Nerve fibers extending along arterioles branch out to envelop renin cell clusters and subsequently continue along efferent arterioles without penetrating into intraglomerular mesangial regions.

**Supplemental Video 5. Renin-expressing cells are extensively distributed in the developing vascular tree at birth, related to Figure 4.**

Three-dimensional structure of renin cells (tdTomato, magenta), renal arteries (αSMA, green), and nuclei (TOPRO3, white) at postnatal day 0 (P0) in a $Ren1^{c\text{-}tdTomato/+}$ mouse kidney.

**Supplemental Video 6. Renin distribution becomes localized prominently in juxtaglomerular regions at P5, related to Figure 4.**

Visualization of renin cells (tdTomato, magenta), arterial tree (αSMA, green), and nuclei (TOPRO3, white) in kidney cortex of a postnatal day 5 (P5) $Ren1^{c\text{-}tdTomato/+}$ mouse.

**Supplemental Video 7. Nerve fibers precede vascular maturation in outer cortex at P0, related to Figure 5.**

Three-dimensional structure illustrating nerve fibers (TUBB3, yellow), arterial walls (αSMA, green), and renin cells (tdTomato, magenta) at postnatal day 0 (P0) in a *Ren1*$^{c\text{-}tdTomato/+}$ mouse kidney.

**Supplemental Video 8. FoxD1-derived vasculature and renin cells at early development, related to Figure 6.**

Continuous cross-sectional visualization of FoxD1-lineage (tdTomato, red) and renin-expressing cells (YFP, yellow) at postnatal day 5 (P5) in a *FoxD1*$^{GC}$*; R26R*$^{tdTomato}$*; Ren1*$^{c\text{-}YFP}$ mouse kidney.

**Supplemental Video 9. Chronic RAS deficiency induces concentric hypertrophy and hyperinnervation, related to Figure 7.**

Three-dimensional imaging of renin cells (tdTomato, magenta), vascular smooth muscle cells (αSMA, green), and nerve fibers (TUBB3, yellow) in a 7-month-old *Ren1*$^{c\text{-}tdTomato/tdTomato}$ mouse kidney.

**Supplemental Video 10. Advanced hypertrophy of renin cells and smooth muscle cells and extensive hyperinnervation narrow afferent arteriolar lumen, related to Figure 7.**

Continuous cross-sectional views depicting relationships among renin cells (tdTomato, magenta), additional layers of smooth muscle (αSMA, green), and nerve fibers (TUBB3, yellow) in afferent arterioles of a 7-month-old *Ren1*$^{c\text{-}tdTomato/tdTomato}$ mouse kidney.

### Supplemental Methods

#### Three-dimensional image data analysis

##### *Length of Renin clusters*

This analysis utilized 3D images captured at 0.36× zoom with a 20× objective lens. For accurate measurements, only AAs within the vascular tree, fully traceable from the bifurcation of the arcuate artery to the glomerular inlet, were included. Renin cluster length was specifically defined as the contiguous region of renin expression extending along AAs upstream from the glomerulus, excluding renin-expressing cells scattered in striped patterns along arterioles. To measure renin cluster lengths associated with Figure 2C, 2D, and Supplementary Figure 1, we first generated 3D renin surface objects from tdTomato signals using the Imaris Surface function. Renin cluster lengths were defined as the continuous linear distance of the renin-expressing region extending proximally from the glomerulus. To quantify these lengths, linear distances were measured between the vertex closest to the glomerulus and the opposite vertex located proximally along the AA for each renin surface object adjacent to the glomerulus. Renin positive lengths were defined as the total length encompassing all regions of renin expression along the AA, including discontinuous or striped expression patterns extending proximally beyond the continuous renin cluster. These lengths were quantified by measuring the linear distance between the vertex closest to the glomerulus of the renin surface object and the most upstream vertex of the renin-positive segments along the arteriole. All measurements were performed using the Measurement Points function of Imaris software.

##### *Volume and number of renin cells*

This analysis utilized higher-resolution 3D images captured at 1.2× zoom with a 20× objective lens, focusing exclusively on cortical AAs. To quantify the number of renin cells related to Figure 2E–2H, we combined the Imaris Surface and Spot functions. Initially, 3D renin cluster surface objects were created from tdTomato signals using the Imaris Surface function. Subsequently, nucleoid spots (diameter 4 μm) were generated from the TOPRO3 nuclear staining signals using the Spot function. Only nucleoid spots located within -0.1 µm of the renin cluster surface objects (i.e., fully contained within renin cell clusters) were counted. The volume of each renin cell cluster was quantified using the Surface function. Renin cluster lengths were measured as described above using the Measurement Points function.

#### *Types of renin cell distribution*

The localization patterns of renin cells along AAs were classified into seven distinct types, based on previously established criteria(1):

Type 1: Renin cells continuously cover the entire length of the AA.

Type 2: Renin cell clusters extend proximally from the glomerulus but do not cover the full length of the arteriole. Clusters ≥60 µm in length were specifically classified as Type 2 (extended pattern), as 85.1% of renin clusters were <60 µm under normal conditions.

Type 3: Renin cells exhibit a striped distribution pattern along the AA.

Type 4: Renin cells are restricted exclusively to the JG region, forming clusters <60 µm.

Type 5: No detectable renin immunoreactivity along the AA.

Type 3/2: A mixed pattern, showing characteristics of both Type 3 and Type 2 distributions within the same arteriole.

Type 3/4: A mixed pattern, showing characteristics of both Type 3 and Type 4 distributions within the same arteriole.

#### *Segmentations of arterial trees*

The segmentations of arterial trees were performed using a combination of Gradient Boosted Tree (GBT) pixel classifier and intensity-based thresholding. Briefly, the fluorescence image in the arterial tree channel was first smoothed using a two-pixel Gaussian filter followed by manual annotation at a single pixel level. GBT algorithm was implemented to train the pixel classifier to distinguish between the arterial tree and tissue autofluorescence. Iterative training and annotations were followed by manual examination. After removing tissue autofluorescence, the arterial tree segmentation was performed by subtracting local background and intensity-based thresholding. The final segmentation results were confirmed by two researchers independently.

**Transmission electron microscopy (TEM)**

For TEM analyses, 2-month-old C57BL/6 mice were subjected to a captopril (0.5 g/L in the drinking water) and low sodium (0.1% Na+) diet for 1 week to facilitate the detection of renin cells. Mice were perfused by cardiac perfusion through the left ventricle with 2% PFA/ 2.5% glutaraldehyde fixation solution. Tissue preparation and imaging were performed at the Molecular Electron Microscopy Core at the University of Virginia. Briefly, tissues were stained with osmium and embedded in epoxy resin. Ultrathin sections (70 nm thickness) were cut using an ultramicrotome, and mounted onto Pioloform coated copper slot grids. Thick sections (1 µm) were also cut and stained with toluidine blue to identify glomeruli present in the thin sections. The thin sections were then post-stained with uranyl acetate and lead citrate before observation and imaging using a Tecnai F20 transmission electron microscope (FEI, Thermo Fisher Scientific) operated at an accelerating voltage of 120 kV.

**RNA isolation and Real-time RT-PCR analysis**

Quantitative real-time PCR (qPCR) analysis was performed in samples from kidney cortices. We extracted total RNA using TRIzol reagent (Thermo Fisher Scientific) and the RNeasy Mini Kit (Qiagen, Dusseldorf, Germany). Reverse transcription (RT) was performed using oligo(dT) primers and M-MLV Reverse Transcriptase (Promega, Madison, WI) at 42°C for 1 hour according to the manufacturer's instructions. Quantitative real-time PCR was conducted using SYBR Green I (Thermo Fisher Scientific) and a CFX Connect system (Bio-Rad Laboratories). PCR was performed with the following primers; *Ngf,* forward: 5'-GGGAGCGCATCGAGTGAC-3', reverse: 5'-CAAAACTCCACCATGCTGCC-3'; *Ntrk1*, forward: 5'-CAGTGGACGGTAACAGCACA-3', reverse: 5'-AGCCCATCCTCTGGAGCTAA-3'; *Bdnf*, forward: 5'-GGCTGACACTTTTGAGCACGTC-3', reverse: 5'-CTCCAAAGGCACTTGACTGCTG-3'; *Ntrk2*, forward: 5'- CCACGGATGTTGCTGACCAAAG-3', reverse: 5'-GCCAAACTTGGAATGTCTCGCC-3'; *Ntf3*, forward:5'-

CTACTACGGCAACAGAGACGCT-3’, reverse: 5’-GGTGAGGTTCTATTGGCTACCAC-3’; *Ntrk3*, forward: 5’- GTCTGATGCGAGCCCTACACC-3’, reverse: 5’-AGAGAACCACCAGAAGGACGCA-3’; *Rps14,* forward: 5’-AGGACCAAGACCCCTGGA-3’, reverse: 5’-ATCTTCATCCCAGAGCGAGC-3’. The mRNA expressions of *Ngf, Ntrk1, Bdnf, Ntrk2, Ntf3* and *Ntrk3* were normalized to *Rps14* expression. Changes in expression of renin from mice ($Ren1^{c\text{-}tdTomato/tdTomato}$) with renin enzymatic deficiency were determined by the ΔΔCt method and shown as relative expression compared to control samples.

**Declaration of generative AI and AI-assisted technologies in the writing process**

During the preparation of this work the authors used ChatGPT 4.5 in order to polish the language and grammar of the article on May and June 2025. After using this tool/service, the authors reviewed and edited the content as needed and took full responsibility for the content of the publication.
